# Assessing the potential of backscattering as a proxy for phytoplankton carbon biomass

**DOI:** 10.1101/2023.03.28.534581

**Authors:** Camila Serra-Pompei, Anna Hickman, Gregory L. Britten, Stephanie Dutkiewicz

## Abstract

Despite phytoplankton contributing roughly half of the photosynthesis on earth and fueling marine food-webs, field measurements of phytoplankton biomass remain scarce. The particulate backscattering coefficient (*b*_*bp*_) has often been used as an optical proxy to estimate phytoplankton carbon biomass (*C*_*phyto*_). However, total observed *b*_*bp*_ is impacted by phytoplankton size, cell composition, and non-algal particles. The lack of phytoplankton field data has prevented the quantification of uncertainties driven by these factors. Here, we first review and discuss existing *b*_*bp*_ algorithms by applying them to *b*_*bp*_ data from the BGC-Argo array in surface waters (*<*10m). We find a *b*_*bp*_ threshold where estimated *C*_*phyto*_ differs by more than an order of magnitude. Next, we use a global ocean circulation model (the MITgcm Biogeochemical and Optical model) that simulates plankton dynamics and associated inherent optical properties to quantify and understand uncertainties from *b*_*bp*_-based algorithms in surface waters. We do so by developing and calibrating an algorithm to the model. Simulated error-estimations show that *b*_*bp*_-based algorithms overestimate/underestimate *C*_*phyto*_ between 5% and 100% in surface waters, depending on the location and time. This is achieved in the ideal scenario where *C*_*phyto*_ and *b*_*bp*_ are known precisely. This is not the case for algorithms derived from observations, where the largest source of uncertainty is the scarcity of phytoplankton biomass data and related methodological inconsistencies. If these other uncertainties are reduced, the model shows that *b*_*bp*_ could be a relatively good proxy for phytoplankton carbon biomass, with errors close to 20% in most regions.

**Plain Language Summary:** Phytoplankton contribute roughly half of the photosynthesis on earth and fuel fisheries around the globe. Yet, few direct measurements of phytoplankton concentration are available. Frequently, concentrations of phytoplankton are instead estimated using the optical properties of water. Backscattering is one of these optical properties, representing the light being scattered backwards. Previous studies have suggested that backscattering could be a good method to estimate phytoplankton concentration. However, other particles that are present in the ocean also contribute to backscattering. In this paper we examine how well backscattering can be used to estimate phytoplankton. To address this question, we use data from drifting instruments that are spread across the ocean and a computer model that simulates phytoplankton and backscattering over the global oceans. We find that by using backscattering, phytoplankton can be overestimated/underestimated on average by ∼20%. This error differs between regions, and can be larger than 100% at high latitudes. Computer simulations allowed us to quantify spatial and temporal variability in backscattering signal composition, and thereby understand potential errors in inferring phytoplankton with backscattering, which could not have been done before due to the lack of phytoplankton data.

**Key Points:**

- Phytoplankton carbon *b*_*bp*_-based algorithms can differ up to an order of magnitude at low *b*_*bp*_ values.
- An algorithm fitted to a global model output shows biases ranging between 15% and 40% in most regions.
- Most uncertainties are due to the relative contribution of phytoplankton to total *b*_*bp*_.

## 1 Introduction

Phytoplankton drive marine food-webs and play an important role in the global carbon cycle. Despite their importance in marine systems, few field measurements of phytoplankton carbon biomass (*C*_*phyto*_) exist due to difficulties in separating phytoplankton from the rest of the microbial community and other organic particles. To overcome this difficulty, phytoplankton carbon biomass is often estimated using optical proxies. One of these optical proxies is the particulate backscattering coefficient (*b*_*bp*_). The particulate backscattering coefficient is an inherent optical property of particles, and represents the light being scattered backwards. The most common way of estimating *C*_*phyto*_ using *b*_*bp*_ is by setting a simple linear regression *C*_*phyto*_ = *β*_0_ + *β*_1_*b*_*bp*_(*λ*) (where *β*_0_ is the intercept, *β*_1_ the slope and *λ* the wavelength of light). For simplicity, we will use the terms “*b*_*bp*_-based algorithm” to refer to this type of linear regression. This kind of algorithm is often used by the marine ecological and biogeochemical communities to understand phytoplankton dynamics (e.g. Behrenfeld et al., 2017; Britten et al., 2021), fish dynamics (e.g. MacNeil et al., 2015; Cheung et al., 2016), estimate carbon export (Siegel et al., 2014) or estimate net primary production (such as in the Carbon-based Production Model (CbPM) or The Carbon, Absorption, and Fluorescence Euphotic-resolving (CAFE) net primary production model, Westberry et al., 2008; Silsbe et al., 2016). However, the sparsity of direct phytoplankton field observations makes it difficult to determine the potential uncertainties linked to using *b*_*bp*_ as a proxy for *C*_*phyto*_. In this study, we first review existing *b*_*bp*_-based algorithms and examine how they differ from each other. Next, we employ a global coupled optics/ecosystem model to quantify and understand uncertainties in *b*_*bp*_-based algorithms.

Backscattering is not a property unique to phytoplankton: all particles in the ocean, such as heterotrophic bacteria, zooplankton, detritus, minerals and water molecules themselves will scatter light (Stramski et al., 2001, 2004; Morel et al., 2007). Furthermore, *b*_*bp*_ is affected by other factors, such as particle size and cell composition (Loisel et al., 2006; Organelli et al., 2018). Small cells are considerably more abundant than larger cells (Sprules & Barth, 2016), and therefore contribute more to the total backscattering than larger cells (Stramski et al., 2001). Organisms with inorganic cell walls, such as coccolithophores, have a high refractive index and scatter more light than naked cells (Voss et al., 1998). In particular, it has been shown that plated coccolithophores and coccoliths (calcite scales detached from cells) are major contributors to *b*_*bp*_ when blooming (Balch et al., 1996). The fact that so many factors affect measured backscattering leads us to question how good of a proxy *b*_*bp*_ is for phytoplankton carbon.

Current *C*_*phyto*_ *b*_*bp*_ algorithms are derived by using chlorophyll-*b*_*bp*_ relationships from either field samples or satellite remote sensing (Behrenfeld et al., 2005), or by using *C*_*phyto*_-*b*_*bp*_ relationships obtained from field samples (Martinez-Vicente et al., 2013; Graff et al., 2015; Qiu et al., 2021). In general, these algorithms are a simple linear regression between these relationships, even though some more complex version have emerged in recent years (Bellacicco et al., 2019, 2020). These studies show a relatively good correlation between *b*_*bp*_ and *C*_*phyto*_, and Graff et al. (2015) also show that *b*_*bp*_ has a higher coefficient of determination (R^2^) with *C*_*phyto*_ than with chlorophyll (Chl) or any other environmental variable, reinforcing the use of this optical property to estimate phytoplankton carbon biomass. However, a problem between these studies is that field samples of *C*_*phyto*_ are scarce and biased towards low latitudes, raising issues about their general applicability. Furthermore, each study has used different methods and assumptions to estimate *C*_*phyto*_ (see section 3), preventing direct comparison of phytoplankton carbon data and algorithms between studies, and increasing the uncertainties of the parameters from the *C*_*phyto*_-*b*_*bp*_ regression.

There are therefore several levels of uncertainties in the relationship between *b*_*bp*_ and *C*_*phyto*_. Methodological uncertainties can emerge by the different sensors, methods and assumptions used to estimate *C*_*phyto*_ in the field. These uncertainties, together with sampling biases, result in differences across existing algorithms. Other uncertainties come simply from the assumption of using *b*_*bp*_ as a proxy for *C*_*phyto*_, which are difficult to validate due to the lack of *C*_*phyto*_ field data. This scarcity can be overcome by using Chl-*b*_*bp*_ relationships, as done in Behrenfeld et al. (2005). However, by using Chl instead of *C*_*phyto*_, a community-averaged *C*_*phyto*_:*Chl* ratio is implicitly assumed to obtain *C*_*phyto*_ from the *Chl*-*b*_*bp*_ relationship (see section 6 in the SI). This, together with an incomplete understanding of the drivers of the *Chl*-*b*_*bp*_ relationship (Barbieux et al., 2018), prevent a reliable derivation of *C*_*phyto*_. To date, despite the wide applications of these *b*_*bp*_-algorithms in the field, the drivers of the *b*_*bp*_-*C*_*phyto*_ relationship are not yet well understood, and potential uncertainties linked to the use of *b*_*bp*_ as a proxy for *C*_*phyto*_ have not yet been quantified.

Here, we use Bgc-Argo float data, satellite remote sensing data, as well as a global ocean circulation model, to assess the potential of *b*_*bp*_ as a proxy for phytoplankton carbon biomass (*C*_*phyto*_). First, we review existing *C*_*phyto*_ algorithms and apply them to BGC-Argo *b*_*bp*_ data to identify the main differences between algorithm parameters and sources of uncertainties. Next, we use a global ocean ecosystem model (the MITgcm Biogeochemistry and Optical model, referred from now on as MITgcmBgc) that accounts for plankton functional types and associated inherent optical properties to understand the drivers of the *C*_*phyto*_-*b*_*bp*_ and *Chl*-*b*_*bp*_ relationships and quantify associated uncertainties under the ideal scenario where *C*_*phyto*_ is known everywhere and at all times. Here, *b*_*bp*_ and Chl from the model are validated against Argo float data. This approach only looks at the uncertainties linked to the use of *b*_*bp*_ as a proxy for *C*_*phyto*_, and does not consider other methodological uncertainties or sampling biases. Nevertheless, the model provides new insights into the variability of *C*_*phyto*_ and *b*_*bp*_, both emergent properties of the model.

## 2 Methods

### 2.1 BGC-Argo data

We used data from the Biogeochemical-Argo floats array (BGC-Argo, https://biogeochemical-argo.org/). BGC-argo floats provide biogeochemical data from the upper 2000 m of the ocean, surfacing around local noon. Sampling time-frequency varies between mission. We extracted the float data using the Bgc-Argo-Mat Matlab toolbox (Frenzel et al., 2021). We extracted quality controlled Chl and *b*_*bp*_ data (flags ”good” or ”probably-good”, Wong et al., 2021) between 2011 and 2021 from the upper 10 m of the ocean (0.2 m resolution) to be comparable to satellite products. As our interest is open ocean, we removed the data that was in coastal regions as defined in Longhurst provinces (Figure S8 Longhurst, 2010). We end up with a total of 64902 data points (from 315 profiles) that span several biomes of the global oceans, with a sampling bias towards the Southern Ocean (see for example figure 3a).

Chl was obtained as a processed data product from the BGC-Argo array, where Chl is derived from fluorescence. Since the ratio of fluorescence to Chl-a can vary due to several reasons (e.g. phytoplankton types, photoacclimation, non-photochemical quenching), the error can be large, potentially reaching ± 300% (Roesler et al., 2017; Bittig et al., 2019), but can be reduced to a maximum ± 40% by locally sampling Chl and obtaining ratios between chlorophyll fluorescence and Chl. The applied correction for non-photochemical quenching follows the method suggested in Terrats et al. (2020) (which is a variation from Xing et al., 2018) and can be found in https://www.euro-argo.eu/content/download/157287/file/D4.2_v1.0.pdf.

Argo floats measure scattering at 700 nm over a range of angles in a small volume (*<*10 mL) of seawater. Backscattering is subsequently derived. Errors in the backscattering coefficient are at maximum 20% (Bittig et al., 2019). Afterwards, the backscattering coefficient is converted into particulate backscattering coefficient by removing the backscattering from seawater (temperature and salinity dependent, Zhang et al., 2009; Schmechtig et al., 2018). The final particulate backscattering values at 700 nm are provided as a BGC-Argo product. The Argo-derived *b*_*bp*_ may underestimate scattering by sufficiently motile zooplankton that can avoid the sensor or by large zooplankton that can cause spikes in the data (Bishop & Wood, 2008).

### 2.2 Satellite remote sensing data

We use the NASA L-3 (https://oceancolor.gsfc.nasa.gov/l3/) *b*_*bp*_(443) and Chl data from the MODIS-Aqua sensor. The near-surface chorophyll-a concentration algorithm uses an empirical relationship derived from in situ measurements of chl-a and blue-to-green band ratios of in situ remote sensing reflectances (Rrs). The average Chl error relative to field data ranges between 16% and 68% in open ocean waters (optical water types 1 to 5, Moore et al., 2009). Particulate backscattering output is estimated using the Generalized Inherent Optical Properties model (GIOP) (Werdell et al., 2013), with a median percent error of 24% relative to Argo float data (Bisson, Boss, Werdell, Ibrahim, & Behrenfeld, 2021). We used climatological monthly mean outputs with a 4 km resolution.

### 2.3 The MITgcm Biogeochemical and Optical model

The MITgcm Biogeochemical and Optical model (MITgcmBgc) is a global ocean circulation model that simulates plankton functional types. The model has several configurations (see section 1 in the SI), and for this study, inherent optical properties of seawater and particles are also included (Dutkiewicz et al., 2015, 2020). The ecosystem component is embedded into a 1^*◦*^ *×* 1^*◦*^ physical global circulation model (the MITgcm, Marshall et al., 1997) that simulates ocean circulation and mixing, and has been constrained by observations. The model resolves several dissolved and particulate carbon pools (e.g. plankton, detritus, dissolved organic matter, dissolved inorganic carbon) and several nutrients within these pools (nitrogen, phosphorus, silica, iron). Here we briefly describe pertinent components of the the ecological model, the most recent optical implementation, and parameterization of backscattering. We describe the version of the Darwin model model used in this study in section 1 of the SI, and a more in-depth description of equations and optics can be found in (Dutkiewicz et al., 2015). Here, the model accounts for several plankton functional types: pico-phytoplankton (*Prochlorococcus, Synechococcus* and pico-eukaryotes), coccolithophores, diatoms, mixotrophs, diazotrophs, zooplankton and heterotrophic bacteria (Figure S1). Each functional type encompasses several cell sizes (Figures 1 and S1). Size affects physiological rates and predator-prey interactions, where we assume that larger organisms eat smaller ones following a fixed predator-prey size ratio (see Dutkiewicz et al., 2020). Community composition in the model emerges from environmental conditions and interactions between organisms (competition and predation).

**Figure 1.**
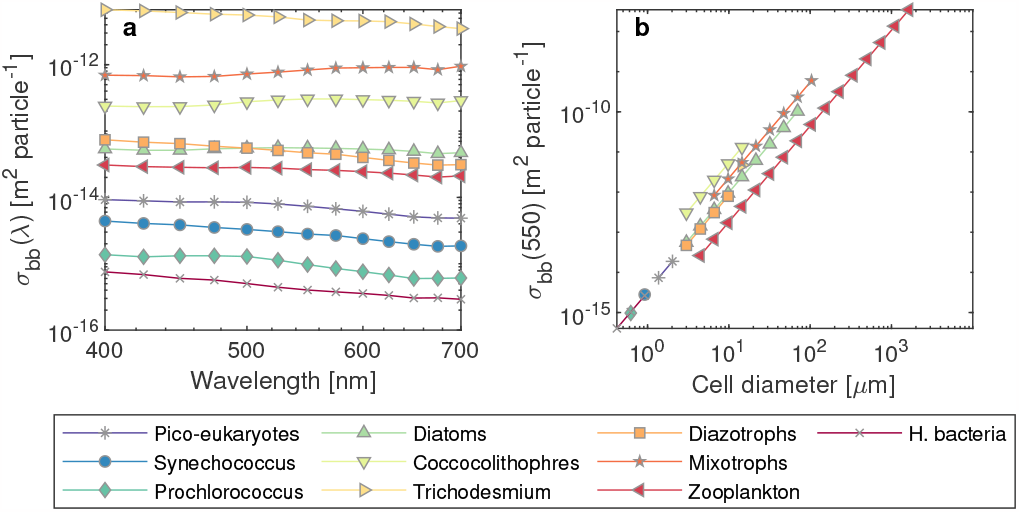
Backscattering cross-sections (*σ*_*bb*_, in m^2^ particle^*−*1^) for each plankton functional type in the MITgcmBgc model against wavelength (a) and plankton body-size (b). The sizes shown in panel (a) are for the smallest organism in each functional group.

#### 2.3.1 Optics in the MITgcmBgc model

Spectral optical properties of water and biology are included in the model (Dutkiewicz et al., 2015, 2018). The model includes a radiative transfer component based on the Ocean–Atmosphere Spectral Irradiance Model (OASIM, Gregg & Casey, 2009), and more fully described in Dutkiewicz et al. (2015). Each type of particle in the water column is represented in the model with its own carbon-specific or Chl-specific absorption, scattering and backscattering. Integrated effects of these optical properties affect the light field available for phytoplankton. As in earlier versions of the model, each phytoplankton functional type and detritus has a specific spectra for absorption and scattering as suggested by observations, and scaled relative to cell size (Dutkiewicz et al., 2015, 2020). New in this latest version, we explicitly include scattering by zooplankton and scattering and absorption by heterotrophic bacteria. Thus, all particles in the model have absorption, scattering and backscattering cross sections associated with them (see figure 1 for backscattering, sections 2-5 in the SI for details on scattering and backscattering, and Dutkiewicz et al. (2015) for absorption).

Here, the total backscattering (*b*_*b*_, in m^*−*1^) is simulated as the sum of the backscattering from water (*b*_*bw*_, in m^*−*1^) and the backscattering by particles (*b*_*bp*_, in m^*−*1^). The particulate backscattering is the sum of the product between the backscattering crosssection (*σ*_*bb*_, in m^2^ mgC^*−*1^) and the total carbon biomass (*C*, mgC m^*−*3^) of each detrital pool or plankton population *i* (of a total of *N*_*p*_ populations). Thus, the total particulate backscattering in the water column is:

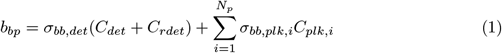

where *σ*_*bb,plk,i*_ and *σ*_*bb,det*_ are the backscattering cross sections of phytoplankton and detritus respectively (in m^2^ mgC^*−*1^), and *C*_*plk,i*_, *C*_*det*_ and *C*_*rdet*_ are the total biomass of each plankton group *i*, labile detritus and refractory detritus in the system respectively (all in mgC m^*−*3^).

Backscattering cross section values were obtained from the literature. For most phytoplankton groups, we use the size-based relation from Vaillancourt et al. (2004) and introduce a scaling factor to differentiate between functional groups. This scaling factor is based on an informed method (see sections 3 and 4 in the SI) to accommodate the species differences by taking backscattering spectra from representative species in culture. Little data is available for zooplankton backscattering, so we assume that they backscatter the same or less than other similar sized unicellular eukaryotes. This assumption is reasonable for unicellular nano-sized zooplankton, which, following a negative size-spectrum slope, will dominate in terms of abundance relative to larger zooplankton, and therefore will also typically dominate the *b*_*bp*_ signal by zooplankton. All the literature sources and derivation of backscattering parameters can be found in the SI.

There are two detrital pools in the Darwin model: an active labile pool and a refractory background pool. The former is important in cycling of carbon and other elements, the latter is introduced in this model for its impact on optics and is crudely parameterized as constant across the globe. There is a high level of uncertainty in *b*_*bp*_ from detritus given the difficulties in both estimating backscattering cross sections of a diverse pool and the parameters needed to convert bulk detrital concentrations (the model variable) to number of particles (needed for the optical impact of this pool). We optimize these uncertain parameters so that model bulk *b*_*bp*_ best matches the BGC-Argo float data (a detailed explanation of this process can be found in section 4 of the SI). We define a constant *q* that combines the conversion from detrital carbon concentration to particle numbers via the size spectrum and particle carbon density assumptions. This combined parameter *q* is optimized. Given the level of uncertainty, we perform a sensitivity analysis of this and other parameters (section 2.4).

The model does not resolve the optical properties of minerals. Minerals can be a major contributors to the *b*_*bp*_ signal (Stramski et al., 2001, 2004). However, our model focuses on the open ocean where the concentration of minerals might be low relative to other optically important constituents. We also implicitly account for particulate inorganic carbon in that we consider a higher backscattering cross-section for coccolithophores (Voss et al., 1998). We do not account for detached coccoliths though. Therefore, the Darwin model likely overestimates somewhat the dependence of *b*_*b*_ on phytoplankton. Thus the error between *C*_*phyto*_ derived using *b*_*bp*_ in the model is likely a lower bound on the error likely found in the real ocean. The model does, however, allow us to investigate the magnitude, variability and sources of the errors.

#### 2.3.2 b_bp_-based C_phyto_ algorithm in the MITgcmBgc model

Following the procedure used for real ocean algorithms (e.g. Behrenfeld et al., 2005; Graff et al., 2015), we calculate model-specific coefficients for a *b*_*bp*_-based algorithm for estimating phytoplankton carbon. The coefficients are found by fitting a linear regression on the linear scale between *C*_*phyto*_ and *b*_*bp*_ and between Chl and *b*_*bp*_. In contrast to the real world, we are in the ideal situation where we know *C*_*phyto*_, Chl and *b*_*bp*_ from the model at all locations; this removes any sampling bias effects and measurement errors.

The values of *C*_*phyto*_ and Chl range over several orders of magnitude. We therefore use a robust regression method to obtain reliable regression parameters at the linear scale. We apply a weighting function to down-weight large outliers in the sum of squares when fitting the regressions and then use Iteratively Reweighted Least Squares (IRLS) to estimate the model parameters. We use the default linear regression MATLAB function with the “robust option” on, which applies a bisquare weighting function to the squared residuals (https://www.mathworks.com/help/stats/robust-regression-reduce-outlier-effects.html).

#### 2.3.3 Algorithm performance assessment

We evaluate the performance of the algorithm derived from the MITgcmBgc model by comparing it against the known modelled phytoplankton carbon. As a measure of algorithm performance we look at the coefficient of determination (R^2^) and the mean absolute error (MAE, as suggested in Seegers et al., 2018; McKinna et al., 2021). The MAE is calculated at the log_10_ scale and afterwards back-transformed to the linear scale:

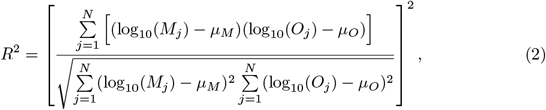

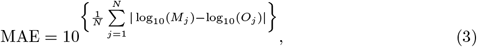

where *N* is the total number of observations, and *O*_*j*_ and *M*_*j*_ are the *j*th “observed” and derived data points (i.e “actual” *C*_*phyto*_ in the Darwin model and the *C*_*phyto*_ derived by the (model) *b*_*bp*_ algorithm respectively). *µ* is the mean of the log-transformed data. The MAE gives a measure of the algorithm absolute bias, that is multiplicative on the back-transformed scale. For example, a MAE of 1.2 means that the algorithm tends to, on average, overestimate/underestimate *C*_*phyto*_ by 20% (in the linear scale).

### 2.4 Sensitivity analysis in the MITgcmBgc

To estimate how parameter uncertainty in the model affect the results of our study, we performed a sensitivity analysis using a Monte Carlo procedure. The uncertain parameters we chose to investigate are the intercept and the slope of the size-scaling relationship of backscattering cross sections for plankton (i.e. intercepts and slopes from figure 1b), and the parameter *q* that encompasses uncertainties for converting detrital concentration to number of particles and implicitly also for the values of the backscattering cross section (section 4 in the SI).

Assuming a uniform probability distribution, we varied the intercepts of *σ*_*bb,plk*_ by ±50% and the slope by ±25%. This combined range covers the values obtained in another study that measured *σ*_*bb*_ for phytoplankton (Whitmire et al., 2010). Given the large uncertainties involved, we varied the parameter *q* over an order of magnitude. We sampled the input space of these parameters using the Latin Hypercube Sampling method.

We performed 500 samples, each with a different value of each input parameter. The sample input matrix was then propagated through the model. The propagation was done offline (i.e. the optics module alone was run on existing model fields), as running the entire model for 500 simulations is computationally unfeasible. The limitation of doing these experiments offline is that there is no feedback between the changes in the inherent optical properties, light trajectories and plankton dynamics, but this method does allows us to efficiently identify the most sensitive optical parameters and explore the sensitivity of our results to these choices.

## 3 Review and further discussion of existing *C*_*phyto*_ *b*_*bp*_-based algorithms

Following development of approaches using backscatter to derive information on phytoplankton and particulate organic carbon (Stramski et al., 1999; Balch et al., 1996; Behrenfeld & Boss, 2003), the use of *b*_*bp*_ as a proxy for *C*_*phyto*_ was presented by Behrenfeld et al. (2005). In that study, the authors argued that even though *b*_*bp*_ is likely more influenced by particles outside the phytoplankton size domain (sub-micron particles), a relation between *b*_*bp*_ and *C*_*phyto*_ can be anticipated, as long as the abundance of these particles co-varies with phytoplankton biomass. Following this assumption, the authors derived an algorithm by subtracting to the *b*_*bp*_ signal a background backscattering value (*b*_*bckg*_) corresponding to a constant stable heterotrophic and detrital components and then by multiplying them by a scalar of 13,000 mgC m^*−*2^. This scalar gave global Chl:*C*_*phyto*_ values close to 0.01 and a *C*_*phyto*_-to-particulate organic carbon (POC) ratio close to 0.3 (average values from the literature). The final equation obtained was *C*_*phyto*_ = 13000(*b*_*bp*_− *b*_*bckg*_). In section 6 of the SI we show that the same equation can be derived by isolating Chl from the linear regression obtained from a *b*_*bp*_-Chl relationship and using an averaged *C*_*P hyto*_:Chl ratio to get *C*_*P hyto*_. The authors argued that Chl:*C*_*phyto*_ values obtained looked reasonable compared to laboratory observations. In later studies, the relationship between *b*_*bp*_ and *C*_*phyto*_ was tested in the field (Martinez-Vicente et al., 2013; Graff et al., 2015; Qiu et al., 2021). These studies showed relatively good correlations between *b*_*bp*_ and *C*_*phyto*_, with R^2^ ranging between 0.53 to 0.7. Graff et al. (2015) also showed that *b*_*bp*_ had a stronger relationship with *C*_*phyto*_ than Chl or any other environmental variable, reinforcing the use of this optical property to estimate phytoplankton carbon biomass.

There are several limitations related to the studies discussed above. First, most field *C*_*phyto*_ and *b*_*bp*_ data is biased towards low latitudes. Second, each study has used different methods and assumptions to estimate in situ *C*_*phyto*_. For instance, Martinez-Vicente et al. (2013) used flow-cytometry and literature cell-mass conversions to obtain the biomass of pico-phytoplankton and some nano-phytoplankton. Qiu et al. (2021) used the same method to estimate pico-phytoplankton biomass and afterwards assumed a size spectrum slope to estimate the biomass for the rest of the phytoplankton community. Graff et al. (2015) used flow cytometry to sort phytoplankton up to cell-sizes of 64 *µ*m, and estimated the carbon content through elemental analysis (Graff et al., 2012). The use of these different methods prevents direct comparison of phytoplankton carbon data and algorithms between studies, and increases the uncertainties of the parameters from the *C*_*phyto*_-*b*_*bp*_ regression.

We plotted all the current algorithms for comparison (Figure 2). After wavelength corrections (see section 7 in the SI), *b*_*bp*_-based algorithms differ remarkably at low values of *b*_*bp*_ and *C*_*phyto*_ (Figure 2): they differ by a factor ∼3 at *b*_*bp*_(470)=10^*−*3^ m^*−*1^, and over an order of magnitude for *b*_*bp*_ values lower 10^*−*3^ m^*−*1^. These discrepancies arise mainly due to differences in the intercepts used in each algorithm. The algorithm from Graff et al. (2015) has the highest intercept, and is the only one to have a positive intercept (*β*_0_ = 0.59, table 1). All the other algorithms have negative intercepts that vary between -4.5 and -22 gC m^*−*3^. On the other hand, algorithms tend to agree at larger value of *b*_*bp*_ and have similar slopes (excluding the algorithm of Martinez-Vicente et al., 2013, which only included pico- and nano-phytoplankton).

**Figure 2.**
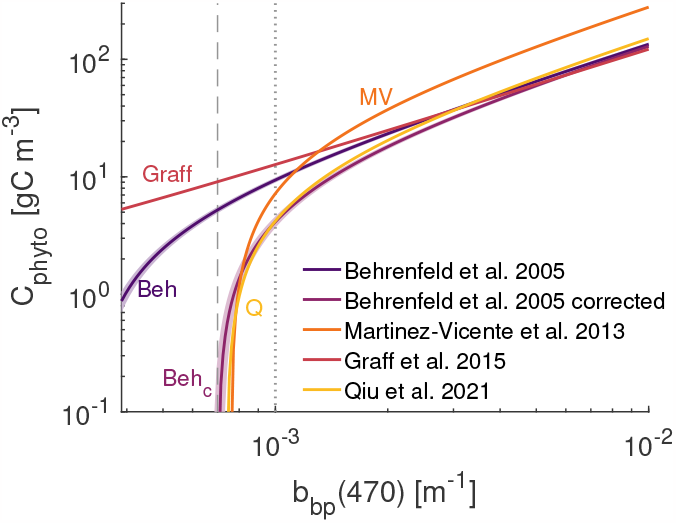
Existing algorithms. All algorithms have been converted to the same wavelength (*λ* = 443) using equation 18 in the SI and a *b*_*bp*_ spectral slope of -1. Shaded areas show the range taken by the algorithms when assuming the *±*0.5 standard deviation of the spectral slope. Dotted line is the *b*_*b,crit*_ threshold and dashed line the *b*_*b,crit*2_ threshold. The “Behrenfeld et al. (2005) corrected” comes from the intercept correction suggested in Qiu et al. (2021). Values of coefficients of these algorithms are listed in table 1.

**Table 1.**
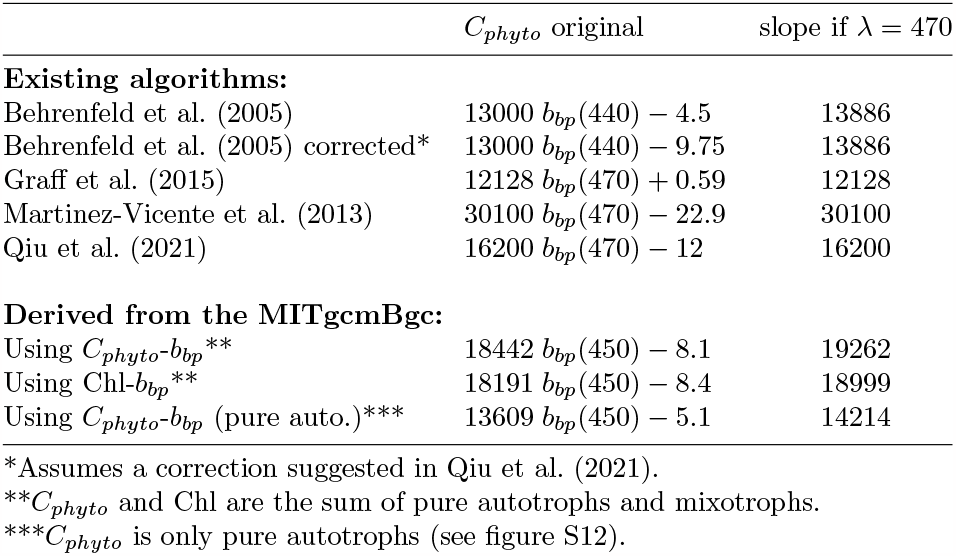
Original algorithms from each study and slopes after wavelength correction to a *λ* = 470 following equations 18 and 19 in the SI.

Using Argo float data, we explore the regions and times of year where *b*_*bp*_ drops below a threshold where the algorithms diverge markedly (*b*_*bp,crit*_(443) = 0.001 m^*−*1^). The *b*_*bp*_ drops below this threshold value in high latitude winters and in some oligotrophic gyres (Figure 3). For oligotrophic regions, most data is below *b*_*bp,crit*_, and about 30% of the observations fall below *b*_*bp*_ values where the algorithms diverge by more than an order of magnitude (*b*_*bp,crit*2_ = 0.0007 m^*−*1^). Temperate regions in winter can have about 60% of their observations below *b*_*bp,crit*_ and 30% below *b*_*bp,crit*2_. Finally, polar regions in winter are always below these two thresholds. Note that the *b*_*bp,crit*_ thresholds relate to the existing algorithms, where the differences below these values emerge out of methodological issues, and probably not due to photoacclimation or differences in the proportion of phytoplankton (we address these later). These are therefore areas where estimated *C*_*phyto*_ differ significantly depending on the *b*_*bp*_ algorithm chosen. More research is needed to constrain *b*_*bp*_ values below *b*_*bp,crit*_.

**Figure 3.**
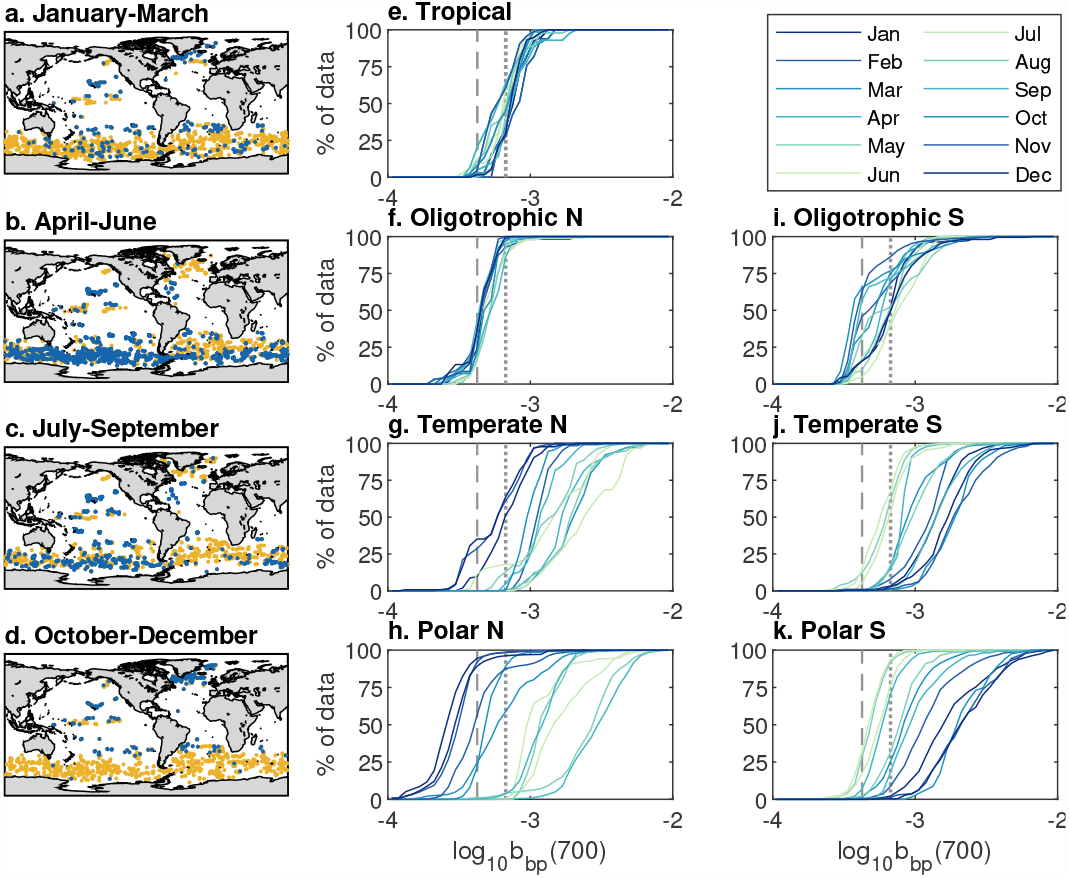
(a-d) location of BGC-Argo data (dots). Data points below the *b*_*b,crit*_ threshold are shown in blue dots. (e-k) cumulative distributions of Argo float *b*_*bp*_ data by biomes in the Northern hemisphere (f-h) and southern hemisphere (i-k). Dotted line is *b*_*b,crit*_ (where algorithms differ by more than a factor of 3), dashed line is *b*_*b,crit*2_ (where algorithms differ by more than an order of magnitude). Both thresholds in this figure have been wave-length corrected using equation 18 in the SI. Note that we only use surface data (*<*10 m), and that coastal areas have been excluded (Figure S10) to be consistent with the output of the model, which does not represent well coastal regions due to its coarse resolution (1^*◦*^ *×* 1^*◦*^).

When following the same approach but for satellite remote sensing (using the MODIS-GIOP sensor and algorithm), most data tends to be above *b*_*bp,crit*_ (Figure S6). For satellite-derived *b*_*bp*_, less than 20% of the data is below *b*_*bp,crit*_ in oligotrophic gyres. This suggests that *b*_*bp*_ derived from satellite (MODIS-GIOP) is overestimated relative to the *b*_*bp*_ derived from Argo floats in regions with low *b*_*bp*_ values. This result is in agreement with results found in other studies, where *b*_*bp*_ derived from different satellite sensors and algorithms were compared with Argo floats data (Bisson et al., 2019; Bisson, Boss, Werdell, Ibrahim, & Behrenfeld, 2021; Bisson, Boss, Werdell, Ibrahim, Frouin, & Behrenfeld, 2021).

Although the availability of in situ data restricts our ability to validate these *C*_*phyto*_ algorithms, we can begin to explore whether the algorithms perform well in certain regions or times of the year by applying the algorithms to the *b*_*bp*_ Argo data and looking at ranges of *C*_*phyto*_ and the Chl:*C*_*phyto*_ (Figures S7 and S8). In this case, the algorithm from Martinez-Vicente et al. (2013) and Qiu et al. (2021) give negative *C*_*phyto*_ values in oligotrophic regions (Figure S7o and S7u). The algorithm from Graff et al. (2015) gives very low Chl:*C*_*phyto*_ ratios in winter in some Polar and sub-polar regions (Figure S8e). These low Chl:*C*_*phyto*_ ratios are characteristic of high light regions, indicating that the Graff et al. (2015) algorithm is probably overestimating *C*_*phyto*_ (possibly due to the high intercept value).

The *C*_*phyto*_ values that these algorithms provide might differ depending on the method used to measure *b*_*bp*_. For instance, satellite remote sensing seems to overestimate *b*_*bp*_ relative to the BGC-Argo measurements (Figure S6). Therefore, if these algorithms were applied to the remote sensing *b*_*bp*_, many of the *C*_*phyto*_ values would not be below *b*_*bp,crit*_, or many of the regions that have negative *C*_*phyto*_ values would probably be positive. We also do not know how the equipment used to measure *b*_*bp*_ in the mentioned field studies compare with remote sensing or BGC-Argo. Therefore, reconciling approaches to measure/estimate *b*_*bp*_ could decrease uncertainties of *C*_*phyto*_ estimates.

## 4 Exploring algorithm uncertainty using the MITgcmBgc model

We first compare the *b*_*bp*_ and Chl outputs from the global ecosystem model (MITgcmBgc) model with the Argo float data (section 4.1). Afterwards, using the MITgcmBgc model output, we quantify the uncertainties of *b*_*bp*_-based algorithms (section 4.3), and explore the potential drivers of these uncertainties (section 4.4). Finally, we evaluate the sensitivity and robustness of our results (section 4.5).

### 4.1 MITgcmBgc model and Argo comparison

We first compare Chl and *b*_*bp*_ from the BGC-Argo and MITgcmBgc model output (Figure 4). Using Bgc-Argo as a reference, the Darwin model is better at simulating *b*_*bp*_ (R^2^ = 0.67, MAE=1.45) than Chl (R^2^ = 0.49, MAE=2.65, Figure 4a,b). The model underestimates Chl by a factor 5 in tropical and some subtropical regions, but follows relatively well the trend in the rest of regions (Figure 4a). The underestimation of Chl in tropical and subtropical regions could be due to the model not representing photoacclimation correctly in these regions (we use Geider, 1987), or due to the coarse resolution of the model, which does not capture sub-mesoscale dynamics that result in the input of nutrients in these less productive regions (see e.g. Clayton et al., 2017; Gupta et al., 2022). Regarding *b*_*bp*_, the Darwin model overestimates by less than a factor of 2 in temperate, sub-polar and polar regions, and slightly underestimates at low latitudes (Figure 4b).

**Figure 4.**
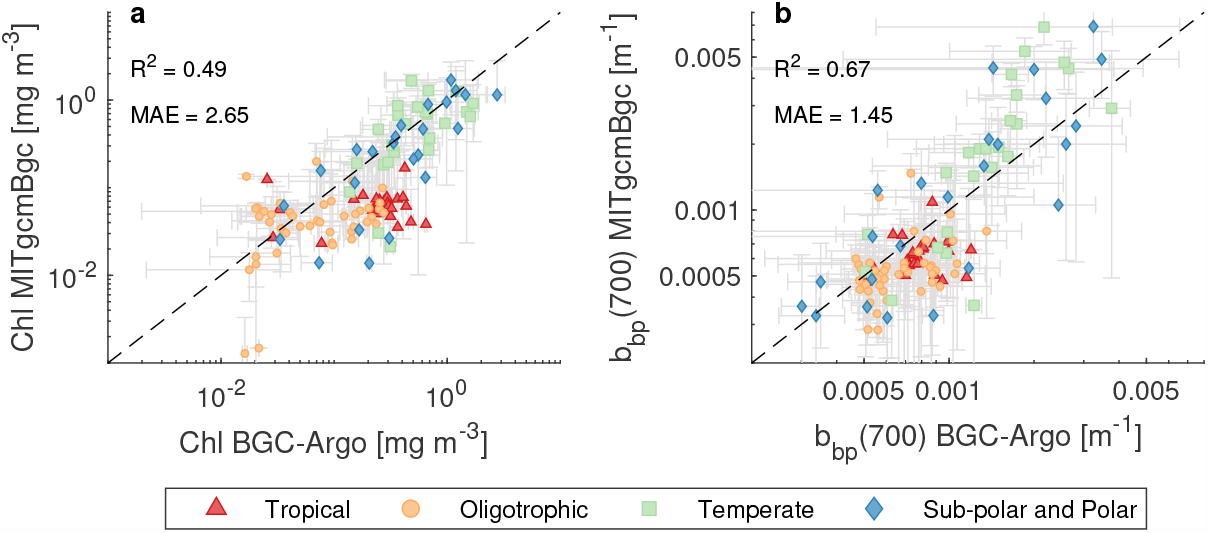
Comparison between the MITgcmBgc model and BGC-Argo surface (*<*10 m) Chl (a) and *b*_*bp*_ (b). Each marker is a monthly average for each biome where there was Argo float data, and grey error bars are the standard deviations. First, bins from the MITgcmBgc were matched to Argo float data-points using a nearest neighbour approach. Afterwards, the data was averaged by month and by biomes, where each biome was defined by grouping Longhurst regions (as seen in figure S10). R^2^ was estimated in the log scale, and MAE was estimated in the log scale and back-transformed afterwards, as shown in equation 3. Dashed line is the 1:1 line.

### 4.2 *b*_*bp*_-based algorithms obtained in the MITgcmBgc model

Since the paucity of real ocean data has prevented a systematic estimation of how well *b*_*bp*_ predicts *C*_*phyto*_, we use the MITgcmBgc model to generate a *C*_*phyto*_-*b*_*bp*_ algorithm (i.e. a linear regression, Figure 5) and test it on the *C*_*phyto*_ from the model. Since current algorithms are generated either by using *C*_*phyto*_ or Chl, we generate two algorithms: one using a *C*_*phyto*_-*b*_*bp*_ relationship (Figure 5a) and another using a [*Chl* × *Q*]-*b*_*bp*_ relationship, where *Q* is a scaling factor that gives reasonable *C*_*phyto*_ values (Figure 5b, the reasoning is similar to the one followed in Behrenfeld et al. (2005) and is described in section 6 of the SI). Each linear regression is fitted to all the surface *C*_*phyto*_ (*<*10 m), Chl and *b*_*bp*_ output data of the global ecosystem model.

**Figure 5.**
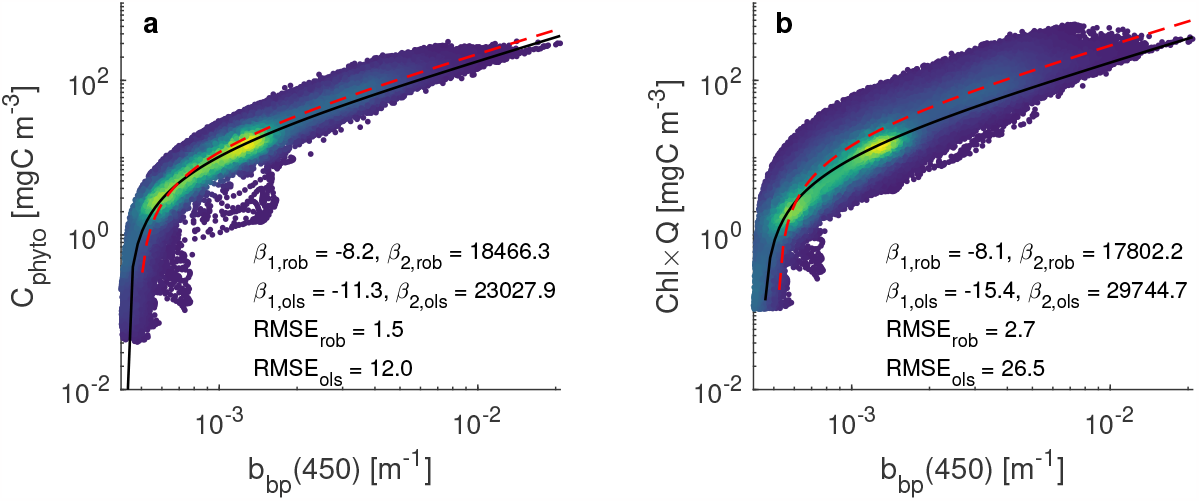
Linear regression of the *C*_*phyto*_-*b*_*bp*_ relationship (a) and the Chl-*C*_*phyto*_ relationship in the MITgcmBgc model for surface waters (*<*10 m). *Q* = 110 and represents a communityaveraged C:Chl ratio as explained in section 6 of the SI. Black continuous line denotes the regression line using the robust regression method (see section 2.3.2), where *β*_1,*rob*_ is the intercept and *β*_2,*rob*_ is the slope. Red dashed line shows the regression line using ordinary least squares (OLS). *C*_*phyto*_ is the summed biomass of pure autotrophs and mixotrophs. Colors show normalized data density. Each dot represents a 1 degree bin of the surface ocean in the model. Chl values below 0.001 mg m^*−*3^ have been removed, as this threshold is close to the detection threshold of the BGC-Argo floats.

Regression parameter values obtained from the model are shown in table 1. We obtain negative intercepts in all cases. Slope values tend to be higher than most algorithms when considering *C*_*phyto*_ and Chl to be the sum of pure autotrophs and mixotrophs, whereas the slope is lower and closer to the ones obtained in Behrenfeld et al. (2005) and Graff et al. (2015) when considering *C*_*phyto*_ to be the biomass of pure autotrophs alone (see also figure S12 in the SI). This could suggest that (in the model) the proportion of mixotrophs relative to pure autotrophs increases in more productive systems. Since mixotrophs contribute to Chl-a, carbon and NPP, from now on we will consider *C*_*phyto*_ as the biomass of both pure autotrophs and mixotrophs.

### 4.3 Error-estimation of the *b*_*bp*_-based algorithms in the MITgcmBgc model

We calibrated two *b*_*bp*_-based *C*_*phyto*_ algorithms to all the surface bins of the MITgcmBgc model (Figure 5). The first model uses *C*_*phyto*_ (Figure 5a), whereas the second model uses *Q* × Chl (Figure 5b), which is an equivalent to *C*_*phyto*_ following the discussion in section 6 of the SI. The regressions performed better when using a robust method rather than ordinary least squares regression method, where the root mean squared errors (RMSE) of the robust method were lower for both models (Figure 5). The spread was wider in the *Q* × Chl-*b*_*bp*_ model, probably due to the use of an averaged community *C*_*phyto*_:Chl ratio (*Q*, units [mgC mgChl^*−*1^]). Note that using this constant factor still allows obtaining variable Chl:C ratios derived from backscattering (Figure S13). In this study, we tried to use a scaling factor *Q* that gave values close to the one of *C*_*phyto*_ in the model. This values is however unknown in the real world, and variations in this parameter can result in substantial overestimation/underestimations of *C*_*phyto*_. Therefore, the overall performance of *C*_*phyto*_ algorithms that use a Chl-*b*_*bp*_ regression might vary depending on the assumed scaling factor *Q*.

Next, we compare month-to-month predicted *C*_*phyto*_ from the algorithm compared to the modeled *C*_*phyto*_ (Figure 6). This pair-wise comparison shows that the *b*_*bp*_-algorithm is able to capture the large scale *C*_*phyto*_ patterns, with R^2^ *>* 0.9 at the global scale (Figure 6e-i). The global monthly MAE (eq. 3) ranges from ∼1.20 to 1.33 when using the algorithm calibrated with *C*_*phyto*_, and from 1.26 to 1.38 when using the algorithm calibrated with Chl. In other words, a *b*_*bp*_-based algorithm, when applied to the model, can overestimate or underestimate *C*_*phyto*_ by ∼ 20% to ∼ 30% on a global average (on the linear scale).

**Figure 6.**
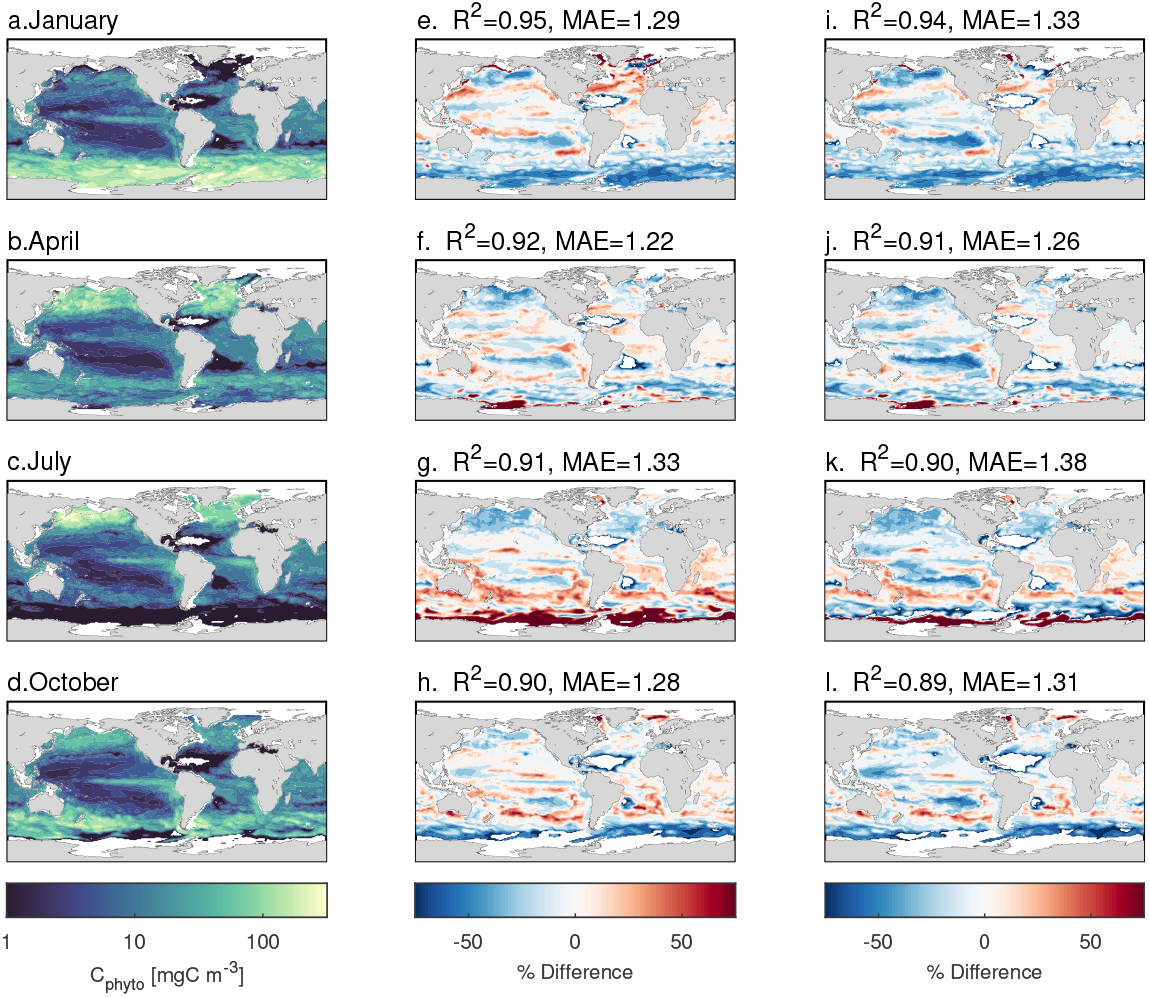
Phytoplankton biomass concentration above 10 m (a-d), and percent difference using the *b*_*bp*_-based algorithm calibrated with *C*_*phyto*_ (e-h) and calibrated with Chl*× Q* (i,l) from figure 5a and b respectively. The shown R^2^ and MAE were estimated in log_10_ scale globally for each month (MAE was backs-transformed to the linear scale as shown in equation 3). White areas represent Chl*<* 0.001 mg m^*−*3^.

The algorithm performance however varies across regions and seasons (Figure 6 and 7). In oligotrophic gyres, errors tend to be below 20% for the algorithm calibrated with *C*_*phyto*_ and below 40% for the algorithm calibrated with Chl (Figure 6 and 7g and h). At higher latitudes, algorithm performance varies seasonally and by ocean bassin. In the sub-polar North Atlantic and North Pacific the algorithm tends to underestimate *C*_*phyto*_ by more than 20% in most regions (Figure 6 and 7a-d). In the Southern Ocean, the algorithm tends to overestimate in winter *>*30 % and underestimate during the rest of the year (Figure 6 and Figure 7j). Overall, the algorithms tend to have errors close to 20 %.

**Figure 7.**
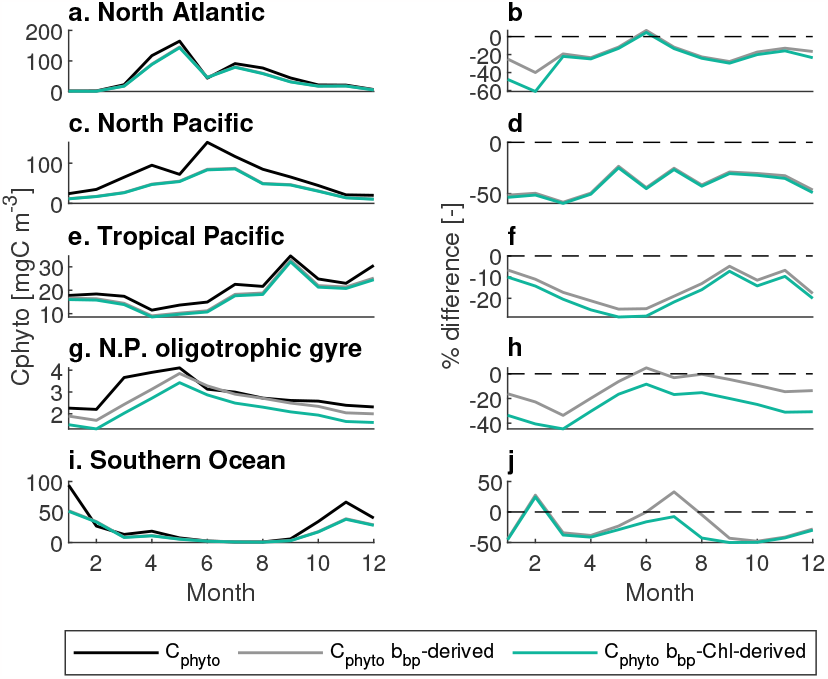
Left-side panels: Seasonal dynamics of phytoplankton biomass (above 10 m) in the MITgcmBgc model (black line), biomass estimated using the *b*_*bp*_-based algorithm using the Cphytp-*b*_*bp*_ relationship (grey lines) and using the Chl-*b*_*bp*_ relationship from figure 5b (green lines) for four regions. Right-side panels: percent difference between *C*_*phyto*_ estimated by the *b*_*bp*_-algorithms relative to the modeled *C*_*phyto*_. Exact locations of each panel are shown in figure S11.

### 4.4 Drivers of the *C*_*phyto*_ *− b*_*bp*_ relationship

To understand what generates the errors and variability in the *C*_*phyto*_ algorithm, we look at the contribution that phytoplankton have on the *b*_*bp*_ signal in the Darwin model (Figure 8). Phytoplankton is the main contributor to *b*_*bp*_ (60%) in spring and summer of seasonal regions. At low latitudes, detrital particles tend to contribute to more than 60% of the signal, whereas phytoplankton mostly account for the rest. Heterotrophic bacteria has a low contribution, except in winter at high latitudes, where it can contribute up to 30% of the *b*_*bp*_ signal. Zooplankton (nano- to meso-) had a negligible contribution to total *b*_*bp*_ (not shown). Also, larger zooplankton could interact with the *b*_*bp*_ signal in different ways that are not captured in the model (e.g. by generating spikes in the *b*_*bp*_ signal due to their size, Bishop & Wood, 2008).

**Figure 8.**
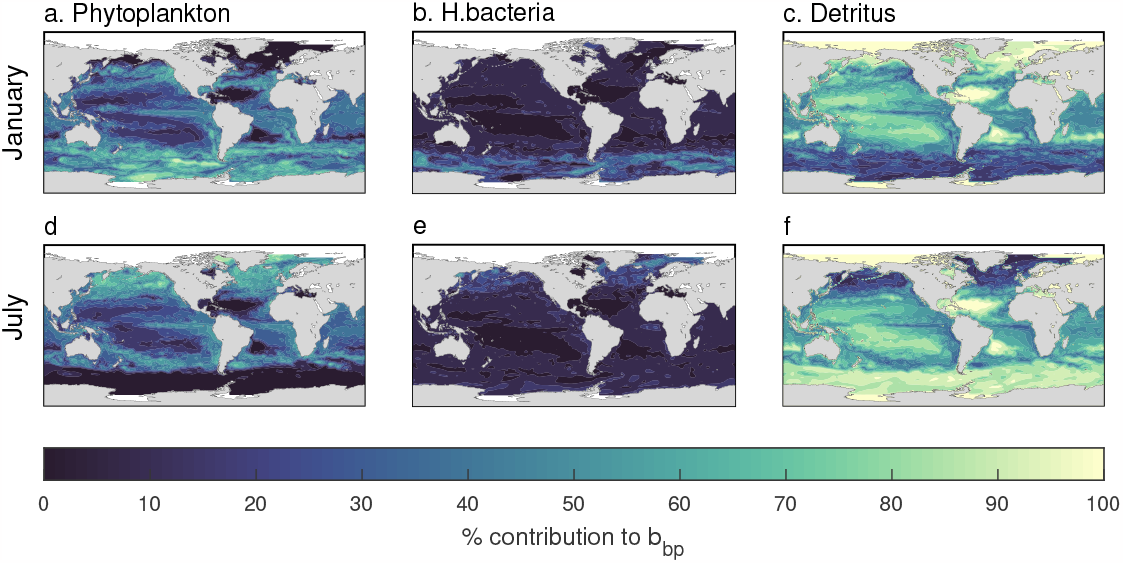
Modeled relative *b*_*bp*_ contribution to the total *b*_*bp*_ signal by phytoplankton (a,d), heterotrophic bacteria (b,e) and detrital particles (c,f) in the surface ocean.

When decomposing the different contributors of the *C*_*phyto*_-*b*_*bp*_ relationship (Figure 9, left-side panels), it can be seen that log_10_(*b*_*bp*_) by phytoplankton alone shows a linear relationship with log_10_(*C*_*phyto*_) (Figure 9a). When adding the effects of heterotrophic bacteria and zooplankton (Figure 9b), a lower *b*_*bp*_ boundary starts to form. However, this lower boundary is much lower than the one set when the contribution by detrital particles is added (Figure 9c). This boundary is higher than most of the *b*_*bp*_ signal set by phytoplankton alone (Figure 9a v.s. 9c). This suggests that regions where *b*_*bp*_ is at its lowest, the signal is mainly dominated by detrital particles.

**Figure 9.**
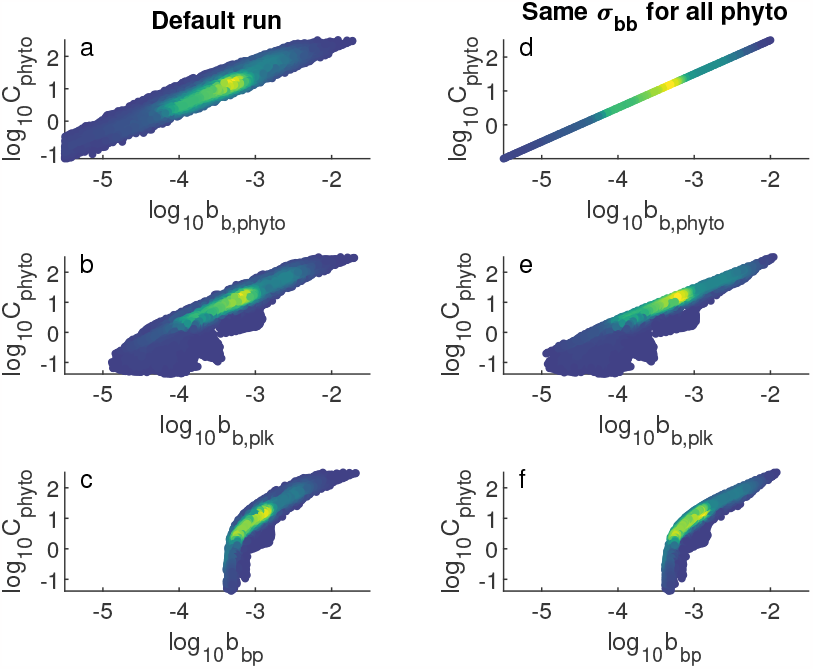
Relationships between *C*_*phyto*_ (mgC m^*−*3^, phytoplankton+mixotrophs) and *b*_*bp*_ (m^*−*1^) from different constituents in the default scenario (a-c) and a scenario where all plankton have the same backscattering cross section (*σ*_*bb*_ = 10^*−*5^, d-f). Backscattering in each panel corresponds to: backscattering by phytoplankton and mixotrophs (a,d), backscattering by plankton (i.e. phytoplankton, mixotrophs, zooplankton and heterotrophic bacteria) only (b,e), and total particulate backscattering (c,f. i.e. all plankton and detritus). Color is the normalized data density.

To understand the effects of phytoplankton functional groups and cell size, we compare a scenario where we assume all phytoplankton have the same backscattering crosssection (Figure 9, right-side panels). This analysis shows that differences in phytoplankton cross-sections do generate some extra variability in the *b*_*bp*_ signal (compare Figure 9a and d). However, at low *b*_*bp*_ values, most of the variability is driven by the contribution of non-algal particles (i.e. detritus, heterotrophic bacteria and zooplankton, Figure 9d vs. e and f).

### 4.5 Sensitivity analysis of the optical parameters in the Darwin model

The sensitivity analysis shows that the mean absolute error of the *C*_*phyto*_ algorithm (MAE from the *C*_*phyto*_-*b*_*bp*_ relationship in figure 6f) can range between 15% and 35% (Figure 10d). The parameter that has the strongest effect on this variation is the slope of the regression between the *b*_*bp*_ cross-section with plankton cell size (Figure 10b, this parameter corresponds to the slope in figure 1b).

**Figure 10.**
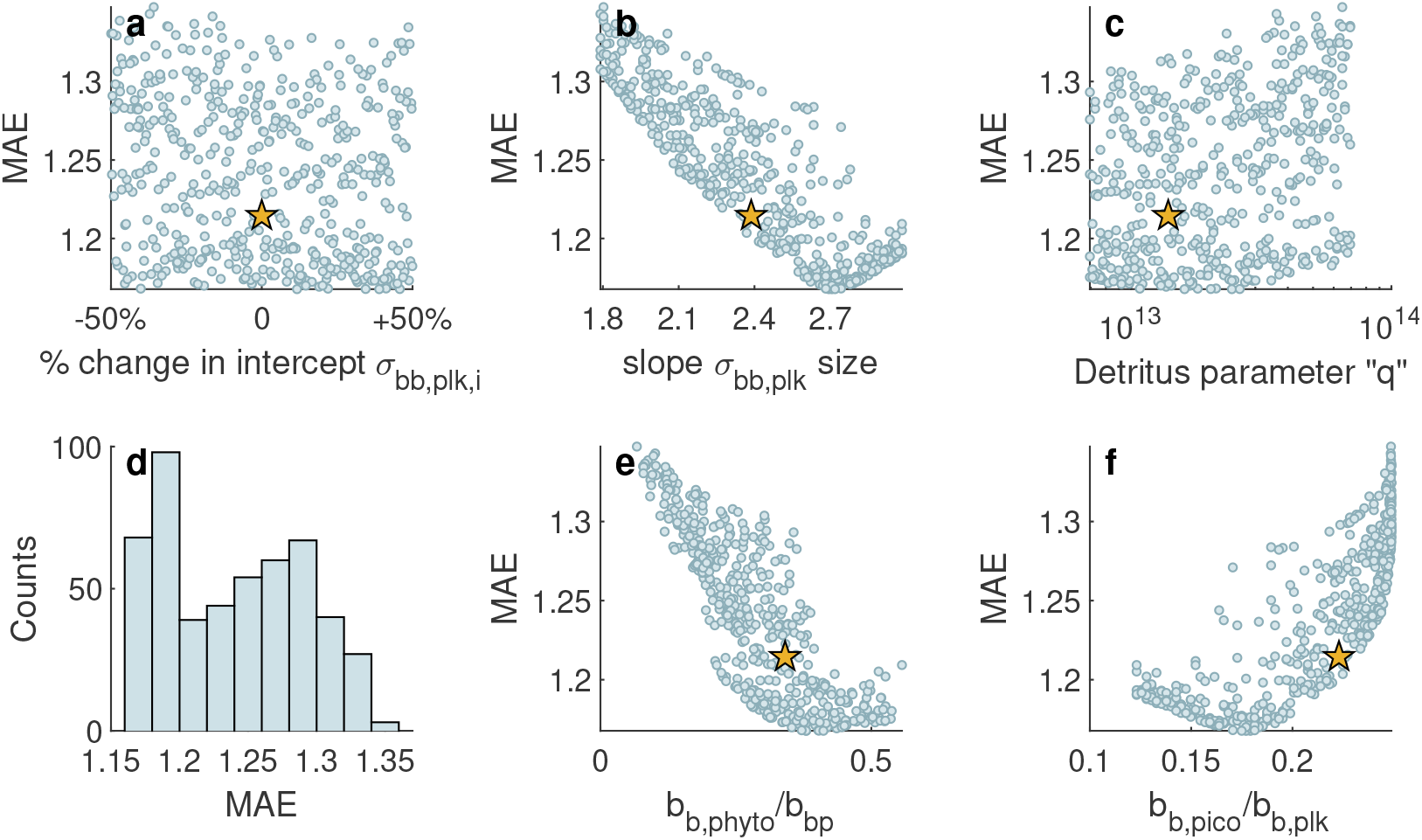
(a-c) Values of MAE for the month of April (corresponding to figure 6f) when randomly varying the three parameters in the sensitivity analysis: intercept and slope of the *σ*_*bb,phyto*_-cell size (corresponding tot he regression parameters in figure 1b), and the *q* parameter that encompasses the uncertainties related to detritus (section 4 in SI). (e) Distribution of MAE from figure 6g after the sensitivity analysis for the three parameters. (e) Variation of MAE with the emergent globally averaged contribution of phytoplankton to total *b*_*bp*_, and with (f) globally averaged contribution of pico-phytoplankton to *b*_*bp*_ by all plankton (*b*_*b,plk*_). Each dot is a run with a random set of parameters. Yellow stars show the values in the default run.

Two potential drivers of the MAE variation are the relative contribution of phytoplankton to total bbp (Figure 10e), and the relative contribution of pico-phytoplankton to the bbp by all plankton (Figure 10f). The slope of the *b*_*bp*_ cross-section with plankton cell size seems to affect these two emergent properties of the model. Lower MAEs occur at steeper slopes, which result in a higher contribution by phytoplankton to total bbp (Figure 10e) and a lower contribution by pico-phytoplankton to the bbp by all plankton (Figure 10f). It is however unclear why there is a kink in MAE with the slope (Figure 10b).

Larger values of the parameter *q* (parameter that encompasses all the uncertainties for the conversion from detritus biomass to number of particles and associated backscattering, section 4 in the SI) resulted in higher MAEs. Larger values of *q* can mean several things: (i) that *σ*_*detr*_ is larger than the one assumed in the default model, (ii) that detrital particles are less dense than we have assumed (affecting our conversion of detrital biomass to number of particles, and therefore resulting in a larger number of detrital particles), and (iii) that the slope of the size spectrum for detritus is steeper than the Junge spectrum assumed. All these factors would increase the contribution of detritus to the overall *b*_*bp*_, reducing the contribution of phytoplankton, and therefore increasing the MAEs.

## 5 Discussion

We have used Bgc-Argo data to identify regions where existing *b*_*bp*_-based *C*_*phyto*_ algorithms differ most in surface waters (*<*10m). Additionally, we have used a global ocean circulation model to assess the magnitude of the potential error in *b*_*bp*_-based algorithms (given perfect knowledge of *C*_*phyto*_ to derive algorithm coefficients) and to understand the drivers of the *b*_*bp*_ signal in the ocean surface.

We show that: (i) there is a threshold of low *b*_*bp*_ where existing algorithms differ up to an order of magnitude. (ii) Regions that are below this threshold are some oligotrophic gyres and high latitudes in winter. Next, in an algorithm calibrated and applied to the Darwin model, we find that (iii) best-case biases in *b*_*bp*_-based algorithms vary markedly across regions and season, ranging from 15% and up to 100%, with most regions having errors close to 20%. Finally, (iv) we show that the variability in the *C*_*phyto*_-*b*_*bp*_ relationship is mainly driven by the varying contribution from non-algal particles (mostly by detritus). In the real world, significant additional uncertainties to algorithms come from the insufficient and non-comparable in-situ measurements of *C*_*phyto*_. As such, our estimates should be thought as best-case biases.

### 5.1 Targeting uncertain regions

We have identified regions where existing algorithms disagree or where *b*_*bp*_-based algorithms might have a poor performance. For example, current algorithms seem to disagree in some oligotrophic regions where the *b*_*bp*_ signal is low, such as around Hawaii. But according to the the global ecosystem model, *b*_*bp*_-algorithms should perform fairly well in this region. Targeting these “high disagreement but high potential” areas could be a first step to reduce uncertainties between current algorithms, as this shows that the uncertainties in existing algorithms are probably driven by other methodological procedures and assumptions not considered in the model (e.g. sensors and approximation to obtain *C*_*phyto*_ from the field).

Winter-time in seasonal regions have several issues: In these regions, currently used algorithms disagree and the model indicates that *b*_*bp*_-based algorithms perform poorly. Obtaining more observations in these regions is also difficult, due to their inaccessibility. However, Chl:*C*_*phyto*_ ratios can help constrain which algorithms perform better. For example, when applying the existing algorithms to the Argo data, it can be seen that the Martinez-Vicente et al. (2013) and Qiu et al. (2021) algorithms give negative *C*_*phyto*_ values in winter of polar and sub-polar regions (Figure S6q,w,s,y), whereas the Graff et al. (2015) algorithm gives suspiciously low Chl:*C*_*phyto*_ ratios in the Northern Polar and sub-polar region (Figure S7e). In these regions, a Chl:*C*_*phyto*_ ratio is expected to be high due to low light levels. Again, in this case the problem is driven by the differences in the intercept, where they seem to be either too low (Martinez-Vicente et al., 2013; Qiu et al., 2021) or too high (Graff et al., 2015).

Even if most algorithms disagree in regions where *b*_*bp*_ values are low, these regions might represent large areas of the ocean (e.g. subtropical gyres). In the MITgcmBgc model, 20% to 40% of the global phytoplankton biomass (depending on the season) is in areas where simulated *b*_*bp*_ is below the *b*_*bp,crit*_ thresholds. Note that the MITgcmBgc model underestimates *b*_*bp*_ in these regions (Figure 4), therefore the total area below this threshold, and therefore the proportion of phytoplankton biomass, might be lower in the real world. Nonetheless, these regions seem to have a considerable role in global *C*_*phyto*_ budgets (and probably NPP budgets) and should not be disregarded.

### 5.2 Contribution of phytoplankton to *b*_*bp*_

Phytoplankton contribution to the bulk *b*_*bp*_ may be larger than previous estimates calculated with Mie theory (Stramski et al., 2001). Using Mie theory, it had been suggested that the main contributors to the *b*_*bp*_ signal are detrital particles (mostly sub-micron sized) and heterotrophic bacteria (Stramski et al., 2001). However, later studies that measured phytoplankton cross-sections from cultures showed that the cross section of phytoplankton were up to an order of magnitude larger than the ones estimated using Mie theory (Vaillancourt et al., 2004; Whitmire et al., 2010). We parameterized our phytoplankton using backscattering cross sections from the latter studies and find that phytoplankton can contribute up to 80% to the bulk *b*_*bp*_ (e.g. during spring blooms). This is in agreement with other studies that suggested that phytoplankton or particles larger than 1 *µ*m could have a significant contribution to the *b*_*bp*_ signal (Dall’Olmo et al., 2009; Brewin et al., 2012; Organelli et al., 2018). Our sensitivity experiments suggest that these assumptions are relatively important, and newer laboratory and theoretical studies to more fully understand the role of plankton versus detrital particles in backscattering are recommended.

### 5.3 Effect of non-algal particles and Chl:C ratios on the Chl-*b*_*bp*_ relationship

Chl-*b*_*bp*_ relationships are often used to understand the relation between *C*_*phyto*_ and *b*_*bp*_ as there is far more Chl data than *C*_*phyto*_. However, it is still somewhat unclear whether non-algal particles or the difference in Chl:C ratios drive the shape of this relationship, especially at low *b*_*bp*_ values (Behrenfeld et al., 2005; Barbieux et al., 2018). From the MITgcmBgc model, it can be seen that the *b*_*bp*_ signal is insensitive to *C*_*phyto*_ or Chl at low values within the upper 10 m of the surface ocean. This is due to the effect of non-algal particles, which override the phytoplankton *b*_*bp*_ signal in oligotrophic regions (setting the intercept of the regressions). These results are partly supported by a study that looked at the drivers of the *b*_*bp*_:Chl relationships using Argo floats (Barbieux et al., 2018). In that study, they found that photoacclimation had practically no effect in the surface layer of the north sub-polar gyre and the Southern Ocean, whereas photoacclimation seemed to affect the *b*_*bp*_:chl ratio for the highest levels of light (*>*0.75 of normalized PAR) in the subtropical gyres and for all PAR levels in the Mediterranean and Black seas (figure 7c in Barbieux et al. (2018)). On the other hand, they did find an important effect of photoacclimation on the *b*_*bp*_:Chl signal when considering deeper layers or the whole mixed layer. In our study, we have not looked at layers below 10 m and both the Mediterranean and Black Seas are not well represented in this version of the MITgcmBgc model due to the coarse resolution. Thus, photoacclimation might play an important role in regions/layers that are not covered in our study. Still, for the regions covered, our finding regarding the effects of non-algal particles are largely in agreement with the ones suggested by Barbieux et al. (2018).

In Behrenfeld et al. (2005), the authors discuss the “bi-linear” trend in the linear scale that they find within the *b*_*bp*_-Chl relationship (i.e. low and high Chl concentrations show different slopes in the linear scale). One of the potential explanations given for this trend are differences in Chl:C ratios. However, we do not find such a clear “bi-linear” trend (in the linear scale) in the Darwin model or in satellite remote sensing output when using the GIOP algorithm with the MODIS-Aqua sensor (Figure S8, noting we used climatological monthly data). We believe that the bi-linear trend in their study might be driven by the algorithm used to estimate *b*_*bp*_ (Garver-Siegel-Maritorena, GSM, semi-analytical algorithm, Maritorena et al., 2002). This algorithm has been shown to overestimate *b*_*bp*_ at low values relative to Argo float data (Bisson et al., 2019), and might not be due to effects of photoacclimation. Nonetheless, whether there is a bi-linear trend or not does not make any difference for the assumptions in their study, and does not change the values of the parameters of their phytoplankton carbon biomass equation.

### 5.4 Errors and uncertainties in *b*_*bp*_-based algorithms

The error we find due to the assumption of *b*_*bp*_ being a good proxy for phytoplankton carbon biomass ranges between 20% and 45% in most regions. This error is of similar magnitude compared to the errors driven by sensor uncertainties and uncertainties related to the approaches to obtain *b*_*bp*_ or Chl. For instance, the backscattering and chlorophyll fluorescence sensors in the BGC-Argo floats have a median error close to 0.11%, and most data showed relative errors lower than 10% (Barbieux et al., 2018). However, errors related to the conversion from fluorescence to Chl increase, reaching up to ±300% (Roesler et al., 2017), and being reduced to ±40% if Chl is sampled locally (Bittig et al., 2019). As for *b*_*bp*_, uncertainties for BGC-Argo are close to 20% (Bittig et al., 2019), and for satellite remote sensing, a bias (calculated as the median ratio of *b*_*bp,sat*_ to *b*_*bp,Argo*_) ranges from 0.77 to 1.6 depending on the algorithm used (Bisson et al., 2019).

The largest source of error for any algorithm likely originates from the way that phytoplankton carbon is derived from field samples or from the assumptions used to derive the scaling factor (here *Q*) used to obtain *C*_*phyto*_ from a Chl-*b*_*bp*_ relationship. Currently a variety of methods are used between field studies (e.g. flow-cytometry and sizespectrum assumptions, or using elemental analysis of carbon and asusmption f Chl:C ratios). Therefore, a standardized method to measure phytoplankton carbon from the field is desirable.

Ideally, measurements should also specify whether mixotrophic plankton are included or not, as these organisms have been shown to be more common than previously thought (Stoecker et al., 2017). We identify two main issues that can arise regarding mixotrophs. The first one is a methodological issue, where most methods for field observations of *C*_*phyto*_ or Chl cannot distinguish mixotrophic contribution. Thus, mixotrophs are included when using field data of Chl and probably of *C*_*phyto*_. The second issue arises when using a Chl-*b*_*bp*_ relationship to derive *C*_*phyto*_. In this case, an extra scaling coefficient is needed to obtain *C*_*phyto*_ (the *Q* factor in this study, see section 6 in the SI). The meaning of this *Q* factor is loosely defined, but considering the units of this factors (mgC mgChl^*−*1^) it can also be considered as a community-averaged C:Chl ratio. Thus, when using a Chl-*b*_*bp*_ relationship to derive *C*_*phyto*_, whether *C*_*phyto*_ is the biomass of pure autotrophs or of autotrophs and mixotrophs, will require different scaling factors *Q*, since mixotrophs might have different C:Chl ratios compared to pure autotrophs.

Finally, other IOPs, such as the beam attenuation coefficient, have been shown to be better proxies for *C*_*phyto*_ or POC than *b*_*bp*_ (Behrenfeld & Boss, 2003; Boss et al., 2015), and can help reduce uncertainties (though note they also encompass both pure autotrophs and mixotrophs). Transmissometers could be mounted on Argo-floats to obtain values of the beam attenuation coefficient (Bernard et al., 2011), complementing estimations derived using *b*_*bp*_ or helping develop new algorithms from satellites.

## 6 Conclusion

The scarcity of phytoplankton field data and the use of different methods and assumptions to determine *C*_*phyto*_ in situ prevents us from being able to estimate uncertainties in algorithms that aim to quantify phytoplankton carbon biomass. Here, we assessed the performance of *b*_*bp*_-based phytoplankton carbon algorithms and quantified their potential uncertainties in the surface ocean (upper 10 m). We showed that existing algorithms can differ by up to an order of magnitude at low *b*_*bp*_-values. By using a global ocean circulation model, we showed that *b*_*bp*_-based algorithms have a global best-case mean absolute error between 15-30%. The algorithm performance declines when using Chl instead of *C*_*phyto*_ to calibrate the *b*_*bp*_ algorithm. Errors were largest when phytoplankton had less impact on the backscattering than other particles (mainly detritus). These error estimates are made under the assumption that *C*_*phyto*_ is known, and therefore do not include other sources of uncertainty. The largest source of uncertainty of any *b*_*bp*_-based algorithm derived from field data will likely be due to the sparsity of in-situ *C*_*phyto*_ and also to the discrepancies in the methods used to measure this quantity. If these other uncertainties are targeted and reduced, *b*_*bp*_ could potentially be a relatively good proxy for *C*_*phyto*_, with errors close to 20% in most regions (according to our model).

Overall, we have shown that a global ecological model can help quantify uncertainties that are currently impossible to estimate from the available real world data. The results of this study advance our understanding of observation- and model-based optical variability in the ocean and its connection to phytoplankton biomass and chlorophyll concentrations. This approach can help reconsider assumptions of some algorithms, and identify ocean conditions to sample that may best improve future algorithms. Continued work in developing accurate remote sensing algorithms for marine ecosystems will improve our ability to monitor marine ecosystems and their response to global change.

## 7 Open Research

Codes to run the model and generate figures, together with model outputs, are available in Zenodo https://doi.org/10.5281/zenodo.7576886.

The BGC-Argo data were collected and made freely available by the International Argo Program and the national programs that contribute to it (https://argo.ucsd.edu, https://www.ocean-ops.org, https://doi.org/10.17882/42182). The Argo Program is part of the Global Ocean Observing System. BGC-Argo float data was extracted using the BGC-Argo-Mat Matlab toolbox (Frenzel et al., 2021).

Satellite remote sensing data was extracted from NASA Goddard Space Flight Center, Ocean Ecology Laboratory, Ocean Biology Processing Group; (2014): MODIS-Aqua Ocean Color Data; NASA Goddard Space Flight Center, Ocean Ecology Laboratory, Ocean Biology Processing Group. http://dx.doi.org/10.5067/AQUA/MODIS_OC.2014.0

## Acknowledgments

We would like to thank I. Cetinić for constructive discussions, and the reviewers of this study for their valuable feedback. This work builds on prior discussions with participants of the Pools of Carbon in the Ocean (POCO) project led by S. Sathyendranath at Plymouth Marine Laboratory and funded by the European Space Agency. We would also like to thank D. Stramski for providing plankton IOPs data that was used to parameterize the former model configuration and part of the new version. This work was supported by multiple grants from the Simons Foundation: the postdoctoral fellowship in Marine microbial ecology funded CSP (#722859) and GLB (#645921), and SD was funded by the Simons Collaboration on Computational Biogeochemical Modeling of Marine Ecosystems (CBIOMES, grant no. 549931). SD additionally acknowledges funding from NASA (Grant 80NSSC22K0153).

## References

Balch, W. M., Kilpatrick, K. A., Holligan, P., Harbour, D., & Fernandez, E. (1996). The 1991 coccolithophore bloom in the central north atlantic. 2. relating optics to coccolith concentration. Limnology and Oceanography, 41 (8), 1684–1696.

Barbieux, M., Uitz, J., Bricaud, A., Organelli, E., Poteau, A., Schmechtig, C., … others (2018). Assessing the variability in the relationship between the particulate backscattering coefficient and the chlorophyll a concentration from a global biogeochemical-argo database. Journal of Geophysical Research: Oceans, 123 (2), 1229–1250.

Behrenfeld, M. J., & Boss, E. (2003). The beam attenuation to chlorophyll ratio: an optical index of phytoplankton physiology in the surface ocean? Deep Sea Research Part I: Oceanographic Research Papers, 50 (12), 1537–1549.

Behrenfeld, M. J., Boss, E., Siegel, D. A., & Shea, D. M. (2005). Carbon-based ocean productivity and phytoplankton physiology from space. Global biogeochemical cycles, 19 (1).

Behrenfeld, M. J., Hu, Y., O’Malley, R. T., Boss, E. S., Hostetler, C. A., Siegel, D. A., … others (2017). Annual boom–bust cycles of polar phytoplankton biomass revealed by space-based lidar. Nature Geoscience, 10 (2), 118–122.

Bellacicco, M., Cornec, M., Organelli, E., Brewin, R., Neukermans, G., Volpe, G., … others (2019). Global variability of optical backscattering by non-algal particles from a biogeochemical-argo data set. Geophysical Research Letters, 46 (16), 9767–9776.

Bellacicco, M., Pitarch, J., Organelli, E., Martinez-Vicente, V., Volpe, G., & Marullo, S. (2020). Improving the retrieval of carbon-based phytoplankton biomass from satellite ocean colour observations. Remote Sensing, 12 (21), 3640.

Bernard, S., Berthon, J., Bishop, J., Boss, E., Claustre, H., Coatanoan, C., … Ulloa, O. (2011). Bio-optical sensors on argo floats. reports of the international ocean-colour coordinating group.

Bishop, J. K., & Wood, T. (2008). Particulate matter chemistry and dynamics in the twilight zone at vertigo aloha and k2 sites. Deep Sea Research Part I: Oceanographic Research Papers, 55 (12), 1684–1706.

Bisson, K., Boss, E., Werdell, P., Ibrahim, A., & Behrenfeld, M. (2021). Particulate backscattering in the global ocean: A comparison of independent assessments. Geophysical research letters, 48 (2), e2020GL090909.

Bisson, K., Boss, E., Werdell, P. J., Ibrahim, A., Frouin, R., & Behrenfeld, M. (2021). Seasonal bias in global ocean color observations. Applied optics, 60 (23), 6978–6988.

Bisson, K., Boss, E., Westberry, T., & Behrenfeld, M. (2019). Evaluating satellite estimates of particulate backscatter in the global open ocean using autonomous profiling floats. Optics express, 27 (21), 30191–30203.

Bittig, H. C., Maurer, T. L., Plant, J. N., Schmechtig, C., Wong, A. P., Claustre, H., … others (2019). A bgc-argo guide: Planning, deployment, data handling and usage. Frontiers in Marine Science, 6, 502.

Boss, E., Guidi, L., Richardson, M. J., Stemmann, L., Gardner, W., Bishop, J. K., … Sherrell, R. M. (2015). Optical techniques for remote and in-situ characterization of particles pertinent to geotraces. Progress in Oceanography, 133, 43–54.

Brewin, R. J., Dall’Olmo, G., Sathyendranath, S., & Hardman-Mountford, N. J. (2012). Particle backscattering as a function of chlorophyll and phytoplankton size structure in the open-ocean. Optics express, 20 (16), 17632–17652.

Britten, G. L., Padalino, C., Forget, G., & Follows, M. J. (2021). Seasonal photoacclimation in the north pacific transition zone. Global Biogeochemical Cycles, e2022GB007324.

Cheung, W. W., Reygondeau, G., & Frölicher, T. L. (2016). Large benefits to marine fisheries of meeting the 1.5 c global warming target. Science, 354 (6319), 1591–1594.

Clayton, S., Dutkiewicz, S., Jahn, O., Hill, C., Heimbach, P., & Follows, M. J. (2017). Biogeochemical versus ecological consequences of modeled ocean physics. Biogeosciences, 14 (11), 2877–2889.

Dall’Olmo, G., Westberry, T., Behrenfeld, M., Boss, E., & Slade, W. (2009). Significant contribution of large particles to optical backscattering in the open ocean. Biogeosciences, 6 (6), 947–967.

Dutkiewicz, S., Cermeno, P., Jahn, O., Follows, M. J., Hickman, A. E., Taniguchi, D. A., & Ward, B. A. (2020). Dimensions of marine phytoplankton diversity. Biogeosciences, 17 (3), 609–634.

Dutkiewicz, S., Hickman, A., Jahn, O., Gregg, W., Mouw, C., & Follows, M. (2015). Capturing optically important constituents and properties in a marine biogeochemical and ecosystem model. Biogeosciences, 12 (14), 4447–4481.

Dutkiewicz, S., Hickman, A. E., & Jahn, O. (2018). Modelling ocean-colour-derived chlorophyll a. Biogeosciences, 15 (2), 613–630.

Frenzel, H., Sharp, J., Fassbender, A., Buzby, N., Plant, J., Maurer, T., … Gray, A. (2021). Bgc-argo-mat: A matlab toolbox for accessing and visualizing biogeochemical argo data. Zenodo. doi: https://doi.org/10.5281/zenodo.4971318

Geider, R. J. (1987). Light and temperature dependence of the carbon to chlorophyll a ratio in microalgae and cyanobacteria: implications for physiology and growth of phytoplankton. New Phytologist, 1–34.

Graff, J. R., Milligan, A. J., & Behrenfeld, M. J. (2012). The measurement of phytoplankton biomass using flow-cytometric sorting and elemental analysis of carbon. Limnology and Oceanography: Methods, 10 (11), 910–920.

Graff, J. R., Westberry, T. K., Milligan, A. J., Brown, M. B., Dall’Olmo, G., van Dongen-Vogels, V., … Behrenfeld, M. J. (2015). Analytical phytoplankton carbon measurements spanning diverse ecosystems. Deep Sea Research Part I: Oceanographic Research Papers, 102, 16–25.

Gregg, W. W., & Casey, N. W. (2009). Skill assessment of a spectral ocean– atmosphere radiative model. Journal of Marine Systems, 76 (1-2), 49–63.

Gupta, M., Williams, R. G., Lauderdale, J. M., Jahn, O., Hill, C., Dutkiewicz, S., & Follows, M. J. (2022). A nutrient relay sustains subtropical ocean productivity. Proceedings of the National Academy of Sciences, 119 (41), e2206504119.

Loisel, H., Nicolas, J.-M., Sciandra, A., Stramski, D., & Poteau, A. (2006). Spectral dependency of optical backscattering by marine particles from satellite remote sensing of the global ocean. Journal of Geophysical Research: Oceans, 111 (C9).

Longhurst, A. R. (2010). Ecological geography of the sea. Elsevier.

MacNeil, M. A., Graham, N. A., Cinner, J. E., Wilson, S. K., Williams, I. D., Maina, J., … others (2015). Recovery potential of the world’s coral reef fishes. Nature, 520 (7547), 341–344.

Maritorena, S., Siegel, D. A., & Peterson, A. R. (2002). Optimization of a semianalytical ocean color model for global-scale applications. Applied optics, 41 (15), 2705–2714.

Marshall, J., Adcroft, A., Hill, C., Perelman, L., & Heisey, C. (1997). A finite-volume, incompressible navier stokes model for studies of the ocean on parallel computers. Journal of Geophysical Research: Oceans, 102 (C3), 5753–5766.

Martinez-Vicente, V., Dall’Olmo, G., Tarran, G., Boss, E., & Sathyendranath, S. (2013). Optical backscattering is correlated with phytoplankton carbon across the atlantic ocean. Geophysical Research Letters, 40 (6), 1154–1158.

McKinna, L. I., Cetinić, I., & Werdell, P. J. (2021). Development and validation of an empirical ocean color algorithm with uncertainties: A case study with the particulate backscattering coefficient. Journal of Geophysical Research: Oceans, 126 (5), e2021JC017231.

Moore, T. S., Campbell, J. W., & Dowell, M. D. (2009). A class-based approach to characterizing and mapping the uncertainty of the modis ocean chlorophyll product. Remote Sensing of Environment, 113 (11), 2424–2430.

Morel, A., Gentili, B., Claustre, H., Babin, M., Bricaud, A., Ras, J., & Tieche, F. (2007). Optical properties of the “clearest” natural waters. Limnology and oceanography, 52 (1), 217–229.

Organelli, E., Dall’Olmo, G., Brewin, R. J., Tarran, G. A., Boss, E., & Bricaud, A. (2018). The open-ocean missing backscattering is in the structural complexity of particles. Nature Communications, 9 (1), 1–11.

Qiu, G., Xing, X., Boss, E., Yan, X.-H., Ren, R., Xiao, W., & Wang, H. (2021). Relationships between optical backscattering, particulate organic carbon, and phytoplankton carbon in the oligotrophic south china sea basin. Optics Express, 29 (10), 15159–15176.

Roesler, C., Uitz, J., Claustre, H., Boss, E., Xing, X., Organelli, E., … others (2017). Recommendations for obtaining unbiased chlorophyll estimates from in situ chlorophyll fluorometers: A global analysis of wet labs eco sensors. Limnology and Oceanography: Methods, 15 (6), 572–585.

Schmechtig, C., Poteau, A., Claustre, H., D’Ortenzio, F., Dall’Olmo, G., & Boss, E. (2018). Processing bgc-argo particle backscattering at the dac level. version 1.4, 07 march 2018.

Seegers, B. N., Stumpf, R. P., Schaeffer, B. A., Loftin, K. A., & Werdell, P. J. (2018). Performance metrics for the assessment of satellite data products: an ocean color case study. Optics express, 26 (6), 7404–7422.

Siegel, D., Buesseler, K., Doney, S. C., Sailley, S., Behrenfeld, M. J., & Boyd, P. (2014). Global assessment of ocean carbon export by combining satellite observations and food-web models. Global Biogeochemical Cycles, 28 (3), 181–196.

Silsbe, G. M., Behrenfeld, M. J., Halsey, K. H., Milligan, A. J., & Westberry, T. K. (2016). The cafe model: A net production model for global ocean phytoplankton. Global Biogeochemical Cycles, 30 (12), 1756–1777.

Sprules, W. G., & Barth, L. E. (2016). Surfing the biomass size spectrum: some remarks on history, theory, and application. Canadian Journal of Fisheries and Aquatic Sciences, 73 (4), 477–495.

Stoecker, D. K., Hansen, P. J., Caron, D. A., & Mitra, A. (2017). Mixotrophy in the marine plankton. Annu. Rev. Mar. Sci, 9 (1), 311–335.

Stramski, D., Boss, E., Bogucki, D., & Voss, K. J. (2004). The role of seawater constituents in light backscattering in the ocean. Progress in Oceanography, 61 (1), 27–56.

Stramski, D., Bricaud, A., & Morel, A. (2001). Modeling the inherent optical properties of the ocean based on the detailed composition of the planktonic community. Applied Optics, 40 (18), 2929–2945.

Stramski, D., Reynolds, R. A., Kahru, M., & Mitchell, B. G. (1999). Estimation of particulate organic carbon in the ocean from satellite remote sensing. Science, 285 (5425), 239–242.

Terrats, L., Claustre, H., Cornec, M., Mangin, A., & Neukermans, G. (2020). Detection of coccolithophore blooms with biogeochemical-argo floats. Geophysical research letters, 47 (23), e2020GL090559.

Vaillancourt, R. D., Brown, C. W., Guillard, R. R., & Balch, W. M. (2004). Light backscattering properties of marine phytoplankton: relationships to cell size, chemical composition and taxonomy. Journal of plankton research, 26 (2), 191–212.

Voss, K. J., Balch, W. M., & Kilpatrick, K. A. (1998). Scattering and attenuation properties of emiliania huxleyi cells and their detached coccoliths. Limnology and oceanography, 43 (5), 870–876.

Werdell, P. J., Franz, B. A., Bailey, S. W., Feldman, G. C., Boss, E., Brando, V. E., … others (2013). Generalized ocean color inversion model for retrieving marine inherent optical properties. Applied optics, 52 (10), 2019–2037.

Westberry, T., Behrenfeld, M., Siegel, D., & Boss, E. (2008). Carbon-based primary productivity modeling with vertically resolved photoacclimation. Global Biogeochemical Cycles, 22 (2).

Whitmire, A. L., Pegau, W. S., Karp-Boss, L., Boss, E., & Cowles, T. J. (2010). Spectral backscattering properties of marine phytoplankton cultures. Optics Express, 18 (14), 15073–15093.

Wong, A., Keeley, R., Carval, T., et al. (2021). Argo quality control manual for ctd and trajectory data.

Xing, X., Briggs, N., Boss, E., & Claustre, H. (2018). Improved correction for non-photochemical quenching of in situ chlorophyll fluorescence based on a synchronous irradiance profile. Optics express, 26 (19), 24734–24751.

Zhang, X., Hu, L., & He, M.-X. (2009). Scattering by pure seawater: Effect of salinity. Optics express, 17 (7), 5698–5710.

