## Supplemental Information for "Assessing the potential of backscattering as a proxy for phytoplankton carbon biomass"

### Contents of this file

1. Configuration of the MITgcmBgc model
2. Parametrization of scattering for plankton in the Darwin model
3. Parametrization of backscattering for plankton in the Darwin model
4. Scattering and backscattering for detrital particles in the Darwin model
5. Scattering and backscattering of seawater
6. Deriving $C_{phyto}$ from a $b_{bp}$-Chl relationship
7. Wavelength corrections
8. Figures

### 1. Configuration of the MITgcmBgc model

In this study we use the MIT Ocean, Biogeochemical, Ecosystem, Optical model (here referred as MITgcmBgc, and often referred as “The Darwin model”). The MITgcm code is available at https://github.com/darwinproject/darwin3, and documentation and equations https://darwin3.readthedocs.io/en/latest/phys_pkgs/darwin.html. All the files needed to run the model for this project, together with the model output, are available in Zenodo https://doi.org/10.5281/zenodo.7576886.

The model resolves the cycling of C, N, P, Fe, Si, O2 through living, detrital and inorganic pools. It is developed to be flexible in the number and type of plankton that is resolves. It follows a “trait-based” approach as each plankton is assigned a number of traits that define its size, trophic, and nutrient status, thermal responses, and for phytoplankton, light affinity. The combination of traits set the plankton’s uptake and growth parameters. The model can be set up for random assignment of traits (see e.g. Follows et al., 2007; Dutkiewicz et al., 2009), traits largely set by size alone (e.g. Ward et al., 2012), traits focusing on functional type (e.g. Dutkiewicz et al., 2015), or traits focused on trophic strategy (e.g. Ward et al., 2012; Zakem et al., 2018). The version of the model used here follows Follett et al. (2022) where the traits are set by trophic strategy, functional type, and size. Trophic strategy included heterotrophic bacteria, phytoplankton, mixotrophs and grazers. Functional types (within the phytoplankton) include pico-phytoplankton (function is dictated by their high nutrient affinity), calcifiers, nitrogen fixers, and silicifiers. Each of the trophic/functional groups outlined above include a range of size classes (see Fig. S1). The model includes 3 size classes of heterotrophic bacteria, 2 size classes of pico-cyanobacteria, 2 size classes of pico-eukaryotes, 5 size classes of calcifiers, 5 size classes of nitrogen fixers, 9 size classes of silicifers, 8 size classes of mixotrophs, and 16 size classes of zooplankton. Phytoplankton have size resolution spanning from 0.6 µm to 140 µm ESD, zooplankton spanning 4.5 µm to 1,636 µm, and bacteria spanning 0.4 µm to 0.9 µm.

Parameters influencing plankton growth, grazing, and sinking are related to size, with specific differences between the six functional groups following Dutkiewicz, Boyd, and Riebesell (2021) (see Table S1). Phytoplankton growth is limited by multiple nutrients (N, P, Fe, and Si in the case of diatoms), light (following Geider et al., 1998), and temperature (following Anderson et al., 2021). Grazing is parameterized using a Holling II function (Holling, 1972) and is size-specific such that grazers can prey upon plankton 5 to 15 times smaller than themselves, with an optimal size of 10 times smaller (Hansen et al., 1997; Kiørboe, 2016). The emergent size distribution of the simulated plankton populations is strongly controlled both by the rate of supply of limiting nutrients (bottom up) and by grazing (top down) (Ward et al., 2014; Dutkiewicz et al., 2021).

### 2. Parametrization of scattering for plankton in the MITgcmBgc model

Scattering cross section values ($\sigma_b(\lambda)$, in m$^2$ particle$^{-1}$) for each plankton type were obtained from the literature (table S1). We begin by assigning a $\sigma_b(\lambda)$ at the observed size to each plankton group (2.1). Next, to be able to propagate the reference $\sigma_b(\lambda)$ values to the rest of plankton sizes in the model, we use the data from Stramski, Bricaud, and Morel (2001) to get a size scaling of $\sigma_b(\lambda)$ (section 2.2). Finally, we convert the units of $\sigma_b(\lambda)$ to m$^2$ mgC$^{-1}$ to estimate the total scattering plankton biomass (section 2.3). The final values in the model for scattering cross sections are shown in figure S3. The following sections provide a detailed explanation of each step.

#### 2.1. Scattering reference values

Most scattering data was obtained from the Stramski et al. (2001) study, who measured $\sigma_b(\lambda)$ for several plankton species at all wavelengths between 350 and 750 nm (fig. S2). With this data-set, we obtained the reference values for heterotrophic bacteria, *Synechococcus*, *Prochlorococus*, diatoms, coccolithophores, and mixotrophs.

For *Trichodesmium*, Dupouy et al. (2008) provides a chloropyll specific $b^*_{p,chl}(\lambda)$ spectrum in ($m^2$ $mgChl^{-1}$). To convert to carbon units, we assume a C:Chl = 100, following LaRoche and Breitbarth (2005).

We did not find measurements of $\sigma_b(\lambda)$ for unicellular zooplankton. We therefore assume that zooplankton have the same $\sigma_b(\lambda)$ as for the small eukaryotes (size-corrected using equation 1 below). This is a fair assumption considering that the smallest unicellular zooplankton will scatter more relative to larger zooplankton, even though the absence of chlorophyll could reduce the $\sigma_b(\lambda)$. So, even if larger zooplankton (e.g. copepods) have different scattering properties (due, for example, to transparency in their bodies or shape), the small abundance of large zooplankton relative to the smaller unicellular ones will result in negligible contribution of these larger organisms to the total scattering and backscattering signals. Note that we do not resolve the "spikes" in the $b_{bp}$ signal generated by the largest zooplankton as seen in Behrenfeld et al. (2019).

#### 2.2. Scaling of scattering cross section with plankton size

Data from Stramski et al. (2001) shows a bi-linear trend at log scale of $\sigma_b(\lambda)$ (m$^2$ particle$^{-1}$) with the cell diameter (fig. S2a), where the slope changes for larger cells. We fit a power-law function for each of the two parts of the curve for the averaged values within each waveband interval in the model (13 intervals between 400 and 700 nm). The exponents $\beta$ of these regression equations are then used to propagate the $\sigma_b(\lambda)$ from the reference diameter $D_{ref}$ ($\mu$m) to the rest of sizes $D$ within each functional group following:

$$\sigma_b(\lambda) = \sigma_{b,ref}(\lambda)\left(\frac{D}{D_{ref}}\right)^{\beta}. \tag{1}$$

### 2.3. Diameter to carbon mass conversions

To scale these cross sections with the biomass of plankton, we convert the units of $\sigma_b$ (m$^2$ particle$^{-1}$) to units of biomass (m$^2$ mgC$^{-1}$). To do so, we first convert the cell diameter to volume using:

$$V = \frac{\pi}{6}D^3 \tag{2}$$

And from volume to carbon mass we use the conversion factor provided in Montagnes, Berges, Harrison, and Taylor (1994):

$$C = (0.109V^{0.991}) \times 10^{-9}; \tag{3}$$

Where $D$ is diameter ($\mu$m), $V$ volume ($\mu$m$^3$), and $C$ carbon mass in a cell (mgC cell$^{-1}$).

### 3. Parametrization of backscattering for plankton in the MITgcmBgc model

Eariler version of the MITgcmBgc model used the backscattering cross section values ($\sigma_{bb}(\lambda)$, in m$^2$ particle$^{-1}$) provided in Stramski et al. (2001). However, that study used Mie theory to obtain these values, which has since been shown to underestimate $\sigma_{bb}(\lambda)$ of plankton by a factor of 5 to more than an order of magnitude (Stramski et al., 2004; Vaillancourt et al., 2004; Whitmire et al., 2010; Organelli et al., 2018). We therefore now use as a reference the regression equation at $\lambda = 510$ from Vaillancourt et al. (2004) that relates $\sigma_{bb}(510)$ (m$^2$ particle$^{-1}$) to cell diameter $D$ ($\mu$m):

$$\sigma_{bb,Vail}(510) = 4 \times 10^{-15} D^{2.387}. \quad (4)$$

See figure S4a for the extracted data.

To distinguish between plankton functional groups, we add a multiplicative factor to equation 4 (table S2). This factor is obtained by averaging the $\sigma_{bb}(510)$ for each functional group in the Vaillancourt et al. (2004) and Whitmire et al. (2010) data-sets (fig. S5c,d) after normalizing all the $\sigma_{bb}(510)$ to the same size (fig. S5a,b). This averaging shows that Dinoflagellates have on average a $\sigma_{bb}(510)$ twice as high as the the one found using equation 4, diatoms have a lower value, while the rest fall close to the average (fig. S5c). The data provided in Whitmire et al. (2010) shows that only dinoflagellates fall above the average $\sigma_{bb}(422)$ (fig. S5d). We therefore chose a conservative value of a factor 2 higher than the average for the mixotrophs in the MITgcmBgc model (which are representative of dinoflagellates).

For Coccolithophores, we use as a reference value provided in Voss, Balch, and Kilpatrick (1998), which gives a value about 5 times higher than the average in equation 4.

For *Trichodesmium*, we assume that $\sigma_{bb}(\lambda) = \sigma_b(\lambda) \times 0.4\%$ following the references in Mouw, Yoder, and Doney (2012) and Subramaniam, Carpenter, and Falkowski (1999).

For heterotrophic bacteria we used the data provided in Vaillancourt et al. (2004).

There is almost no data of $\sigma_{bb}(\lambda)$ for zooplankton. One study (Basedow et al., 2019) provided values for large and red-pigmented copepods (*Calanus finmarchicus*), which are not so relevant for our model. Another study mentioned that the presence of chlorophyll increased $\sigma_{bb}(\lambda)$ in plankton cells (Vaillancourt et al., 2004). Given this assessement, we chose the $\sigma_{bb}(\lambda)$ of zooplankton to be equal or lower than that of phytoplankton (of the same size). After parameter tuning, we end up with a $\sigma_{bb}(\lambda)$ about 5 times lower than the one values obtained from equation 4.

Finally, to propagate across the rest of wavebands in the model, we assumed that $\sigma_{bb}$ had the same spectral shape as $\sigma_b$. (Whitmire et al., 2010) provided some $\sigma_{bb}$:$\sigma_b$ ratios, that are mostly flat across wavelength, except for wavelengths higher than 650 $\mu$m, where the ratio increased slightly. The spectral shapes of $\sigma_{bb}$ used in this study are shown in figure 1 of the main text, and have been obtained by:

$$\sigma_{bb}(\lambda) = \frac{\sigma_{bb}(510)\sigma_b(\lambda)}{\sigma_b(510)} \tag{5}$$

### 4. Parametrization for detrital particles in the MITgcmBgc model

The scattering and backscattering of detrital particles are the most uncertain parameters of the model. We needed to make several assumptions, and there are few field or laboratory data to validate them. In addition, we also needed to make assumptions to convert the single detrital carbon concentration pool of the MITgcmBgc model to a numbers of detrital particles. To decrease the uncertainties in these conversions, we started by a default run and parameters values as explained in the sections below. Next, we constrained some of these parameter values by comparing the model $b_{bp}$ output to Argo float data. The following sections explain the assumptions made for the default run and the further adjustments made to these parameters.

#### 4.1. Scattering and backscattering for detrital particles

For detrital particles, we use the scattering and backscattering cross sections estimated in Stramski et al. (2001). Despite the uncertainties mentioned in their study, we use these values as a starting point, but then optimize these other parameters related to detritus in order to best match Argo-float data. The equations provided in Stramski et al. (2001) are:

$$\sigma_{b,det}(\lambda) = 0.1425\lambda^{-0.9445} \times 10^{-12} \tag{6}$$

$$\sigma_{bb,det}(\lambda) = 5.881 \times 10^{-4}\lambda^{-0.8997} \times 10^{-12} \tag{7}$$

where $\sigma_{b,det}(\lambda)$ is in m$^2$ particle$^{-1}$ and $\lambda$ in nm. To estimate these cross-sections Stramski et al. (2001) used Mie theory and assumed an assemblage of particles ranging from 0.05 to 500 $\mu$m in diameter. These particles follow a Junge differential size distribution (Junge, 1963) described by a power function with a slope of -4. They also assumed a wavelength-independent real part of refractive index ($n = 1.04$, relative to water).

#### 4.2. Size spectrum of detrital particles

There is only one detrital concentration pool in the MITgcmBgc model (in dimensions of mass per volume). Therefore, to get a number of detrital particles we follow the same assumptions as in Stramski et al. (2001), that is, assuming the detrital pool is made up of particles following a number size spectrum ($N_s$, in particles m$^{-3}$ $\mu$m$^{-1}$) with an exponent of -4 (following a Junge-type size distribution):

$$N_s = \kappa D^{\gamma} \tag{8}$$

where $D$ is the diameter of a paticle (in $\mu$m), $\kappa$ is the coefficient and $\gamma$ the exponent.

To convert the detrital concentration to number of particles, we need to estimate the value of $\kappa$. We do this by converting the previous size spectrum (eq. 8) into a concentration spectrum using the particle carbon mass ($m$, in mgC particle$^{-1}$) and the relation between carbon mass and particle diameter ($m = \alpha D^{\beta}$). We start by assuming that this conversion is the same as for plankton $m = 10^{-9.8} D^{2.82}$ (eq. 3), even thought the value will probably be lower (later discussed). Thus, the new size spectrum (in mgC m$^{-3}$ $\mu$m$^{-1}$) is:

$$B_s = N_s m = \kappa D^{\gamma} m = \kappa D^{\gamma} \alpha D^{\beta} = \kappa \alpha D^{\gamma+\beta} = \kappa \alpha D^{\delta} \tag{9}$$

Hence, $\delta = \gamma + \beta = -1.18$. We solve the integral to get the total detrital carbon mass within the size range $D_{min} = 0.05$ and $D_{max} = 500$ $\mu$m:

$$B = \int_{D_{min}}^{D_{max}} \kappa \alpha D^{\delta} \mathrm{d}D = \kappa \alpha \left[ \frac{D_{max}^{\delta+1}}{\delta + 1} - \frac{D_{min}^{\delta+1}}{\delta + 1} \right] = \kappa \alpha c \tag{10}$$

where $c$ is the second term in the equation above and is constant. Now we get that:

$$\kappa = \frac{B}{\alpha c} \tag{11}$$

We can now get the total number of particles $N$ (# m$^{-3}$) by solving the integral and inserting equation 11 into equation 8. The final equation to get the number of particles from the total biomass is then:

$$N = \int_{D_{min}}^{D_{max}} \kappa D^{\gamma} \mathrm{d}D = B\frac{1}{\alpha c}\left[\frac{D_{max}^{\gamma+1}}{\gamma+1} - \frac{D_{min}^{\gamma+1}}{\gamma+1}\right] = Bq \tag{12}$$

Since all the terms in $q$ are constant, we use it as our conversion factor in the model to obtain the number of particles from the detrital biomass. Following the values we have provided, $q$ is equal to $q = 2.18 \times 10^{12}$. This value has many uncertainties, we therefore constrain it by tuning this parameter to the BGC-Argo data. The new values of $q$ after tuning is provided in the following section.

#### 4.3. Constraining uncertainties in the parameters related to detritus

The parameter $q$ encapsulates the uncertainties of conversion to biomass corrections, but implicitly, also the ones from the scatter and backscattering cross sections for detritus. That is because the term to calculate $b_{bp}$ from detritus biomass is multiplicative: $b_{b,detr} = (C_{det}+C_{rdet})q\sigma_{bb,det}$. To constrain the value of "q", we manually tune this parameter while comparing the total $b_{bp}$ from the MITgcmBgc model with the one from the BGC-Argo float. We start doing this offline. Once we reach a value that we think is reasonable, we run again the whole model with the new value and compare again with the BGC-Argo. We repeated this procedure until we obtained a reasonable fit. After tuning, the new value obtained is $q = 1.39 \times 10^{13}$.

The new $q$ value that best matches observations is slightly higher than the one form the previous model versions. This can mean several things: (i) $\sigma_{detr}$ should be larger than the one assumed in (Stramski et al., 2001) (this is possible given the differences observed in the $\sigma_{plk}$ obtained using Mie theory and the assumptions in (Stramski et al., 2001) to match the bulk $b_{bp}$), (ii) detrital particles are much less dense than we assumed (could be possible due to preferential remineralization of carbon by heterotrophic bacteria), (iii) the slope of the size spectrum for detritus is steeper (on average). Overall, it is probably a combination of all these factors. In our default model simulation we use a $1.39 \times 10^{13}$, though we conduct several sensitivity studies to explore how different values affect our results (see sections 2.4 and 4.4 in the main text).

### 5. Scattering and backscattering of seawater int he MITgcmBgc model

We use the backscattering coefficient of pure seawater ($b_{bsw}$) suggested in Morel et al. (2007). In their study, they assumed $b_{bsw}$ to be half of the scattering of pure seawater ($b_{sw}$ at 20°C), where, $b_{sw}$ was taken as the scattering of pure water from Buiteveld, Hakvoort, and Donze (1994) increased by a factor 1.3 to account for the presence of salt.

### 6. Deriving $C_{phyto}$ from a $b_{bp}$-Chl relationship

To obtain $C_{phyto}$ from a $b_{bp}$-Chl relationship, one simply needs a conversion factor. That is, if we start with the following regression equation:

$$Chl = \alpha_0 + \alpha_1 b_{bp}, \tag{13}$$

where $\alpha_0$ (mgChl m$^{-3}$) is the intercept and $\alpha_1$ (mgChl m$^{-2}$) the slope. To convert Chl to $C_{phyto}$, we simply need a conversion factor ($Q$) of units mgC mgChl$^{-1}$:

$$C_{phyto} = QChl = Q(\alpha_0 + \alpha_1 b_{bp}) = \alpha_{0,C} + \alpha_{1,C} b_{bp}, \tag{14}$$

where $\alpha_{0,C} = Q\alpha_0$ (in mgC m$^{-3}$) and $\alpha_{1,C} = Q\alpha_1$ (in mgC m$^{-2}$). $Q$ can be understood as a community-averaged C:Chl ratio, this is why in our main text we use the term $Chl \times Q$ to get $C_{phyto}$ when we only have Chl information (see section 4.2 and figure 5b in the main text).

This is a similar assumption taken when $C_{phyto}$ is derived using, for example, satellite-derived Chl and $b_{bp}$. The values obtained in the equation used in Behrenfeld, Boss, Siegel, and Shea (2005) ($C_{phyto} = 13000[b_{bp} - 0.00035]$) can be obtained following a similar reasoning. In this equation, 13,000 is a scalar in units of mgC m$^{-2}$ that was chosen to give satellite Chl:C values close to 0.010 and an average phytoplankton contribution to total particulate organic carbon of 30%. The term 0.00035 m$^{-1}$ is a background value estimated using least squares regression analysis of the linear portion of the $b_{bp}$-Chl relationship, that is, the intercept of the regression equation:

$$b_{bp} = b_{bckg} + \beta Chl, \tag{15}$$

where $b_{bckg}$ (m$^{-1}$) is the intercept and $\beta$ (m$^2$ mgChl$^{-1}$) the slope. Isolating Chl from the equation we get:

$$Chl = \frac{1}{\beta}(b_{bp} - b_{bckg}) \tag{16}$$

Now, to get $C_{phyto}$ from this equation, we can use the same conversion factor ($Q$) with units mgC mgChl$^{-1}$. This results in the following:

$$C_{phyto} = QChl = \frac{Q}{\beta}(b_{bp} - b_{bckg}) \tag{17}$$

The term $Q/\beta$ has units of mgC m$^{-2}$, same as the $13,000$ term. Thus, one could argue that for the regression in Behrenfeld et al. (2005) the term $13,000 = Q/\beta$. Given the units of $Q$, this scaling term can be understood as a community-averaged C:Chl ratio.

Assuming that the slope in figure 1 of Behrenfeld et al. (2005) is ∼0.0068, the Q term would be equivalent to 88.4, that is, a Chl:C ratio close to 0.0113. This is in agreement with the reasoning that they make in their study, where they sought to obtain a Chl:C ratio close to 0.010. These are essentially the same assumptions we make in our study, but we start directly by fitting the regression line to the Chl v.s. $b_{bp}$ instead of to the $b_{bp}$ v.s. Chl. Note that having this community-averaged C:Chl ratio does not prevent obtaining variable C:Chl ratios, as those are captured by the variability in data-points (figure S13).

There is however one thing that should be considered when reversing the regression equation (e.g. isolating the Chl as done above). A problem might arise if a type I regression method is performed (as is the least squares regression analysis). Type I regression methods are asymmetric, meaning that different coefficients can be obtained if a regression is fitted to the $x$ v.s. $y$ axis or $y$ v.s. $x$ axis (i.e. Chl v.s. $b_{bp}$ or $b_{bp}$ v.s. Chl). Perhaps a type II regression (symmetric method) should be considered in these cases.

### 7. Wavelength corrections

Depending on the approach used, $b_{bp}$ measurements are performed at different wavelengths ($\lambda$), and therefore, a wavelength correction needs to be performed to compare across algorithms. For instance, Argo floats measure $b_{bp}$ at $\lambda = 700$, whereas the $b_{bp}$ provided by NASA from the GIOP model in MODIS-Aqua is at $\lambda = 443$. The field studies that have looked at the $C_{phyto}$-$b_{bp}$ relationship often provide their $b_{bp}$ at $\lambda = 470$. To be able to compare across algorithms (fig. 2 in main text) the effect of the different wavelength needs to be corrected. Assuming that $b_{bp}$ at any wavelength is:

$$b_{bp}(\lambda) = b_{bp}(\lambda_0)\left(\frac{\lambda}{\lambda_0}\right)^{\gamma} = b_{bp}(\lambda_0)\delta(\lambda_0, \lambda) \tag{18}$$

where $\lambda_0$ is the reference wavelength and $\gamma$ the spectral slope of $b_{bp}$; we can correct the wavelength of $b_{bp}$ in the algorithms by introducing the correction factor $\delta(\lambda_0, \lambda)$. For example, assuming the $C_{phyto}$-$b_{bp}$ relationship, the wavelength correction in the regression equation becomes:

$$C_{phyto} = \beta_0 + \beta_1 b_{bp}(\lambda_0) = \beta_0 + \frac{\beta_1}{\delta(\lambda_0, \lambda)} b_{bp}(\lambda) \quad (19)$$

where $\beta_0$ and $\beta_1$ are the intercept and slope respectively. This essentially results in a new slope $\beta_{1,new} = \beta_1/\delta(\lambda_0, \lambda)$ (right-side column of table 1 in the main text).

The spectral slope ($\gamma$) of $b_{bp}$ ranges between -0.5 to -2 for regions dominated by organic particles (Loisel et al., 2006; Reynolds et al., 2016). We chose $\gamma = -1 \pm 0.5$, which is close to the mean observed slope (Loisel et al., 2006; Reynolds et al., 2016). When including the standard deviation values, we show that the effect of using different $\gamma$ within reasonable values is rather low (fig. 2 in main text). Finally, the MITgcmBgc model simulates $b_{bp}$ for wavelengths between 400 and 700 nm in 13 intervals (where each value quoted is the centre of the each waveband), and therefore no wavelength correction is needed.

### References

Anderson, S., Barton, A., Clayton, S., Dutkiewicz, S., & Rynearson, T. (2021). Marine phytoplankton functional types exhibit diverse responses to thermal change. *Nature communications*, *12*(1), 1–9.

Basedow, S. L., McKee, D., Lefering, I., Gislason, A., Daase, M., Trudnowska, E., … Falk-Petersen, S. (2019). Remote sensing of zooplankton swarms. *Scientific reports*, *9*(1), 1–10.

Behrenfeld, M. J., Boss, E., Siegel, D. A., & Shea, D. M. (2005). Carbon-based ocean productivity and phytoplankton physiology from space. *Global biogeochemical cycles*, *19*(1).

Behrenfeld, M. J., Gaube, P., Della Penna, A., O'malley, R. T., Burt, W. J., Hu, Y., … others (2019). Global satellite-observed daily vertical migrations of ocean animals. *Nature*, *576*(7786), 257–261.

Buiteveld, H., Hakvoort, J., & Donze, M. (1994). Optical properties of pure water. In *Ocean optics xii* (Vol. 2258, pp. 174–183).

Dupouy, C., Neveux, J., Dirberg, G., Rottgers, R., Tenório, M. B., & Ouillon, S. (2008). Bio-optical properties of the marine cyanobacteria trichodesmium spp. *Journal of Applied Remote Sensing*, *2*(1), 023503.

Dutkiewicz, S., Boyd, P. W., & Riebesell, U. (2021). Exploring biogeochemical and ecological redundancy in phytoplankton communities in the global ocean. *Global change biology*, *27*(6), 1196–1213.

Dutkiewicz, S., Follows, M. J., & Bragg, J. G. (2009). Modeling the coupling of ocean ecology and biogeochemistry. *Global Biogeochemical Cycles*, *23*(4).

Dutkiewicz, S., Hickman, A., Jahn, O., Gregg, W., Mouw, C., & Follows, M. (2015). Capturing optically important constituents and properties in a marine biogeochemical and ecosystem model. *Biogeosciences*, *12*(14), 4447–4481.

Follett, C. L., Dutkiewicz, S., Ribalet, F., Zakem, E., Caron, D., Armbrust, E. V., & Follows, M. J. (2022). Trophic interactions with heterotrophic bacteria limit the range of prochlorococcus. *Proceedings of the National Academy of Sciences*, *119*(2), e2110993118.

Follows, M. J., Dutkiewicz, S., Grant, S., & Chisholm, S. W. (2007). Emergent biogeography of microbial communities in a model ocean. *science*, *315*(5820), 1843–1846.

Geider, R. J., MacIntyre, H. L., & Kana, T. M. (1998). A dynamic regulatory model of phytoplanktonic acclimation to light, nutrients, and temperature. *Limnology and oceanography*, *43*(4), 679–694.

Graff, J. R., Westberry, T. K., Milligan, A. J., Brown, M. B., Dall'Olmo, G., van Dongen-Vogels, V., ... Behrenfeld, M. J. (2015). Analytical phytoplankton carbon measurements spanning diverse ecosystems. *Deep Sea Research Part I: Oceanographic Research Papers*, *102*, 16–25.

Hansen, P. J., Bjørnsen, P. K., & Hansen, B. W. (1997). Zooplankton grazing and growth: Scaling within the 2-2,-$\mu$m body size range. *Limnology and oceanography*, *42*(4), 687–704.

Holling, C. (1972). Ecological models: a status report.

Junge, C. E. (1963). Air: Chemistry and radioactivity. *Int Geophys. Ser.*

Kiørboe, T. (2016). Foraging mode and prey size spectra of suspension-feeding copepods and other zooplankton. *Marine Ecology Progress Series*, *558*, 15–20.

LaRoche, J., & Breitbarth, E. (2005). Importance of the diazotrophs as a source of new nitrogen in the ocean. *Journal of Sea Research*, *53*(1-2), 67–91.

Loisel, H., Nicolas, J.-M., Sciandra, A., Stramski, D., & Poteau, A. (2006). Spectral dependency of optical backscattering by marine particles from satellite remote sensing of the global ocean. *Journal of Geophysical Research: Oceans*, *111*(C9).

Martinez-Vicente, V., Dall'Olmo, G., Tarran, G., Boss, E., & Sathyendranath, S. (2013). Optical backscattering is correlated with phytoplankton carbon across the atlantic ocean. *Geophysical Research Letters*, *40*(6), 1154–1158.

Montagnes, D. J., Berges, J. A., Harrison, P. J., & Taylor, F. (1994). Estimating carbon, nitrogen, protein, and chlorophyll a from volume in marine phytoplankton. *Limnology and Oceanography*, *39*(5), 1044–1060.

Morel, A., Gentili, B., Claustre, H., Babin, M., Bricaud, A., Ras, J., & Tieche, F. (2007). Optical properties of the "clearest" natural waters. *Limnology and oceanography*, *52*(1), 217–229.

Mouw, C. B., Yoder, J. A., & Doney, S. C. (2012). Impact of phytoplankton community size on a linked global ocean optical and ecosystem model. *Journal of Marine Systems*, *89*(1), 61–75.

Organelli, E., Dall'Olmo, G., Brewin, R. J., Tarran, G. A., Boss, E., & Bricaud, A. (2018). The open-ocean missing backscattering is in the structural complexity of particles. *Nature Communications*, *9*(1), 1–11.

Qiu, G., Xing, X., Boss, E., Yan, X.-H., Ren, R., Xiao, W., & Wang, H. (2021). Relationships between optical backscattering, particulate organic carbon, and phytoplankton carbon in the oligotrophic south china sea basin. *Optics Express*, *29*(10), 15159–

15176.

Reynolds, R. A., Stramski, D., & Neukermans, G. (2016). Optical backscattering by particles in arctic seawater and relationships to particle mass concentration, size distribution, and bulk composition. *Limnology and Oceanography*, *61*(5), 1869–1890.

Stramski, D., Boss, E., Bogucki, D., & Voss, K. J. (2004). The role of seawater constituents in light backscattering in the ocean. *Progress in Oceanography*, *61*(1), 27–56.

Stramski, D., Bricaud, A., & Morel, A. (2001). Modeling the inherent optical properties of the ocean based on the detailed composition of the planktonic community. *Applied Optics*, *40*(18), 2929–2945.

Subramaniam, A., Carpenter, E. J., & Falkowski, P. G. (1999). Bio-optical properties of the marine diazotrophic cyanobacteria trichodesmium spp. ii. a reflectance model for remote sensing. *Limnology and Oceanography*, *44*(3), 618–627.

Vaillancourt, R. D., Brown, C. W., Guillard, R. R., & Balch, W. M. (2004). Light backscattering properties of marine phytoplankton: relationships to cell size, chemical composition and taxonomy. *Journal of plankton research*, *26*(2), 191–212.

Voss, K. J., Balch, W. M., & Kilpatrick, K. A. (1998). Scattering and attenuation properties of emiliania huxleyi cells and their detached coccoliths. *Limnology and oceanography*, *43*(5), 870–876.

Ward, B. A., Dutkiewicz, S., & Follows, M. J. (2014). Modelling spatial and temporal patterns in size-structured marine plankton communities: top–down and bottom–up controls. *Journal of Plankton Research*, *36*(1), 31–47.

Ward, B. A., Dutkiewicz, S., Jahn, O., & Follows, M. J. (2012). A size-structured foodweb model for the global ocean. *Limnology and Oceanography*, *57*(6), 1877–1891.

Whitmire, A. L., Pegau, W. S., Karp-Boss, L., Boss, E., & Cowles, T. J. (2010). Spectral backscattering properties of marine phytoplankton cultures. *Optics Express*, *18*(14), 15073–15093.

Zakem, E. J., Al-Haj, A., Church, M. J., van Dijken, G. L., Dutkiewicz, S., Foster, S. Q., ... Follows, M. J. (2018). Ecological control of nitrite in the upper ocean. *Nature communications*, *9*(1), 1–13.

**Table S1.** Sources of the scattering cross sections $\sigma_{bb}$.

| Model Analogue Name | Experimental Species | Reference |
|---|---|---|
| Prochlorococcus | Generic Prochlorophyte PROC | Stramski et al. (2001) |
| Synechococcus | Generic Synechococcus | Stramski et al. (2001) - original data from Morel et al. 1993 |
| Pico-Eukaryote | *Micromonas pusilla* | DuRand et al. 2002 |
| Coccolithophores | *Emiliania huxleyi* | Stramski et al. (2001) - original data from Ahn et al. 1992 |
| Diazotroph (unicellular) | Same as for Synechococcus | Same as for Synechococcus |
| Trichodesmium | *Trichodesmium erythraeum* | Dupouy et al. 2008 |
| Pico-Eukaryotes | *Isochrysis galbana* | Stramski et al. (2001) - original data from Ahn et al. 1992 |
| Diatom | *Chaetoceros curvisetus* | Stramski et al. (2001) - original data from Bricaud et al. 1988 |
| Mixotrophic dinoflagellate | *Prorocentrum micans* | Stramski et al. (2001) - original data from Ahn et al. 1992 |
| Zooplankton | Same as for pico-eukaryotes (size-scaled) | |
| Heterotrophic bacteria | multi species assemblage | Stramski et al. (2001) - original data from Stramski and Kiefer 1990 |

[a] Footnote text here.

**Table S2.** Reference backscattering cross sections $\sigma_{bb}$ ($m^2$ particle$^{-1}$) for each plankton functional type in the MITgcmBgc model.

| **Plankton type** | **Value assumed** | **Comment and sources** |
|---|---|---|
| Pico-Eukaryotes | $\sigma_{bb,vail}(510)$ | Eq. 4, obtained from Vaillancourt et al. (2004) |
| Prochlorococcus | $\sigma_{bb,vail}(510)$ | Eq. 4, obtained from Vaillancourt et al. (2004) |
| Synechococcus | $\sigma_{bb,vail}(510)$ | Eq. 4, obtained from Vaillancourt et al. (2004) |
| Diazotrophs | $\sigma_{bb,vail}(510)$ | Eq. 4, obtained from Vaillancourt et al. (2004) |
| Diatoms | $\sigma_{bb,vail}(510)$ | Eq. 4, obtained from Vaillancourt et al. (2004) |
| Mixotrophs | $\sigma_{bb,vail}(510) \times 2$ | Eq. 4 and scaling factor approximated from fig. S5c,d, obtained from Vaillancourt et al. (2004) |
| Heterotrophic bacteria | $\sigma_{bb,vail}(510)$ | Eq. 4, obtained from Vaillancourt et al. (2004) |
| Coccolithophores | $\sigma_{bb,vail}(510) \times 5$ | Eq. 4, obtained from Vaillancourt et al. (2004) and scaling factor approximated from the $\sigma_{bb}$ values in Voss et al. (1998) |
| Zooplankton | $\sigma_{bb,vail}(510)/5$ | Eq. 4, obtained from Vaillancourt et al. (2004) and tunes scaling factor |
| Trichodesmium | $b(\lambda) \times 0.004$ | Value obtained from Mouw et al. (2012) and Subramaniam et al. (1999) |

[a] Footnote text here.

### 8. Additional figures

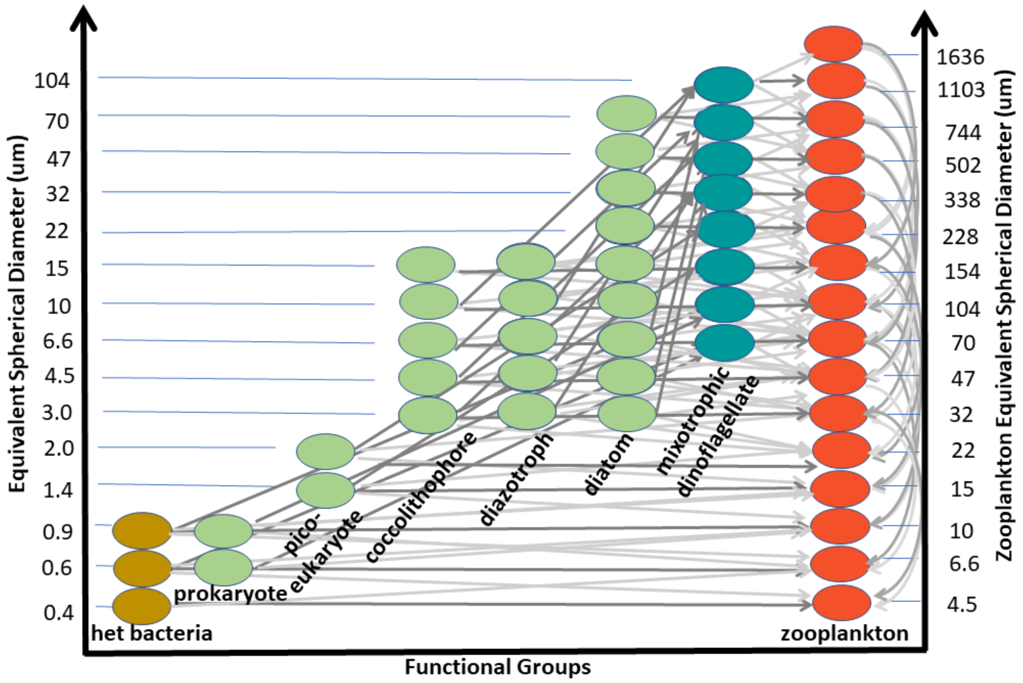


**Figure S1.** Schematic of the ecological component of the MITgcmBgc model version used in this study showing size and functional groups. Left y-axis shows size for heterotrophic bacteria, phytoplankton, and mixotrophs. Right-side y-axis shows size for zooplankton. Grey lines show who is eaten by who, with thicker lines corresponding to preferred grazing size, and thinner lines grazing at a lower preference.

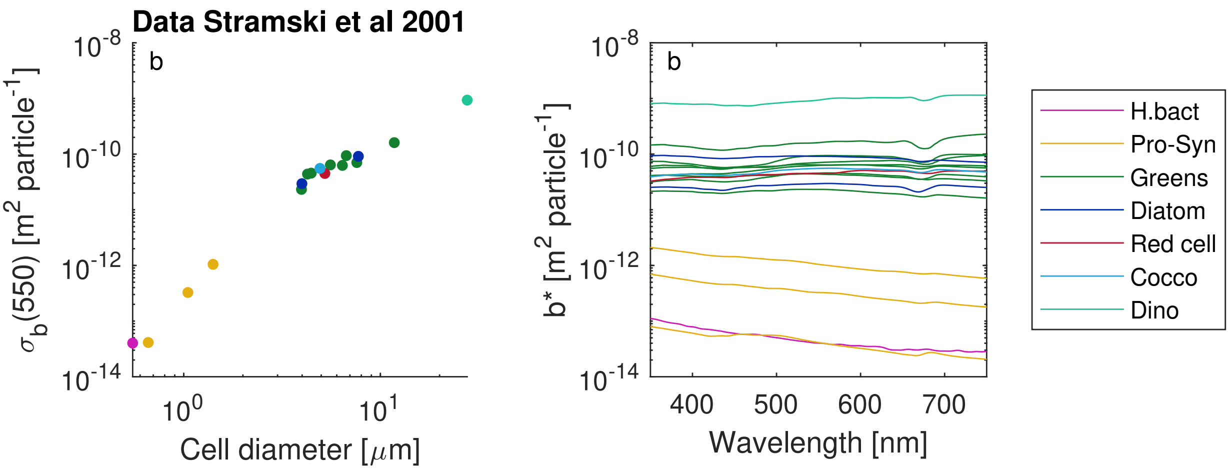


**Figure S2.** Total scattering cross section data from Stramski et al. (2001).

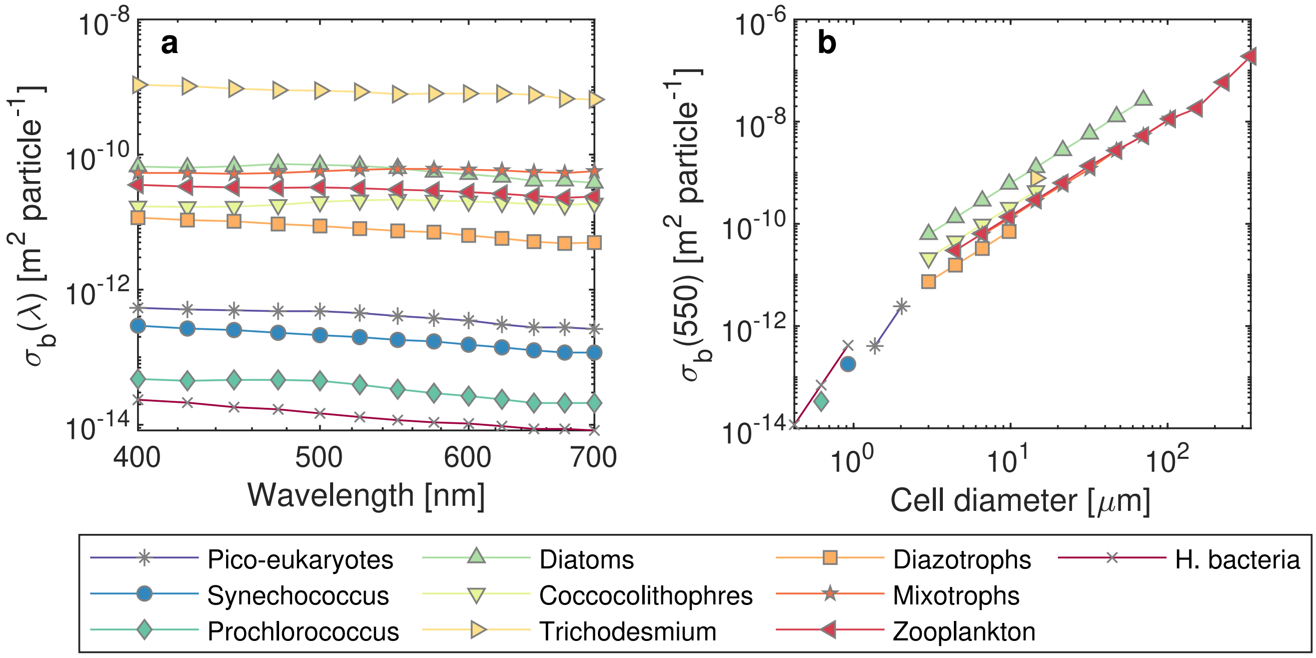


**Figure S3.** Total scattering cross section in the MITgcmBgc model.

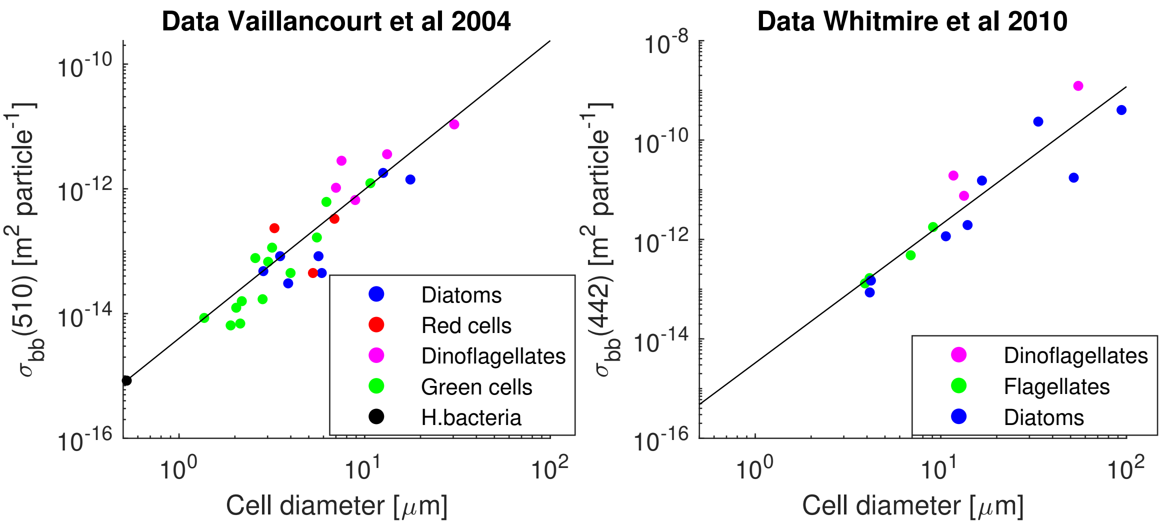


**Figure S4.** Data extracted from Vaillancourt et al. (2004) and Whitmire et al. (2010) grouped by functional groups (e.g. diatoms, dinoflagellates and heterotrophic bacteria) or by similar traits (all green or red flagellated cells that are not part of the previous functional groups mentioned). Black lines are the fit to the data: $\sigma_{bb,vail}(510) = 4 \times 10^{-15} D^{2.39}$ and $\sigma_{bb,whit}(442) = 10^{-14.48} D^{2.78}$.

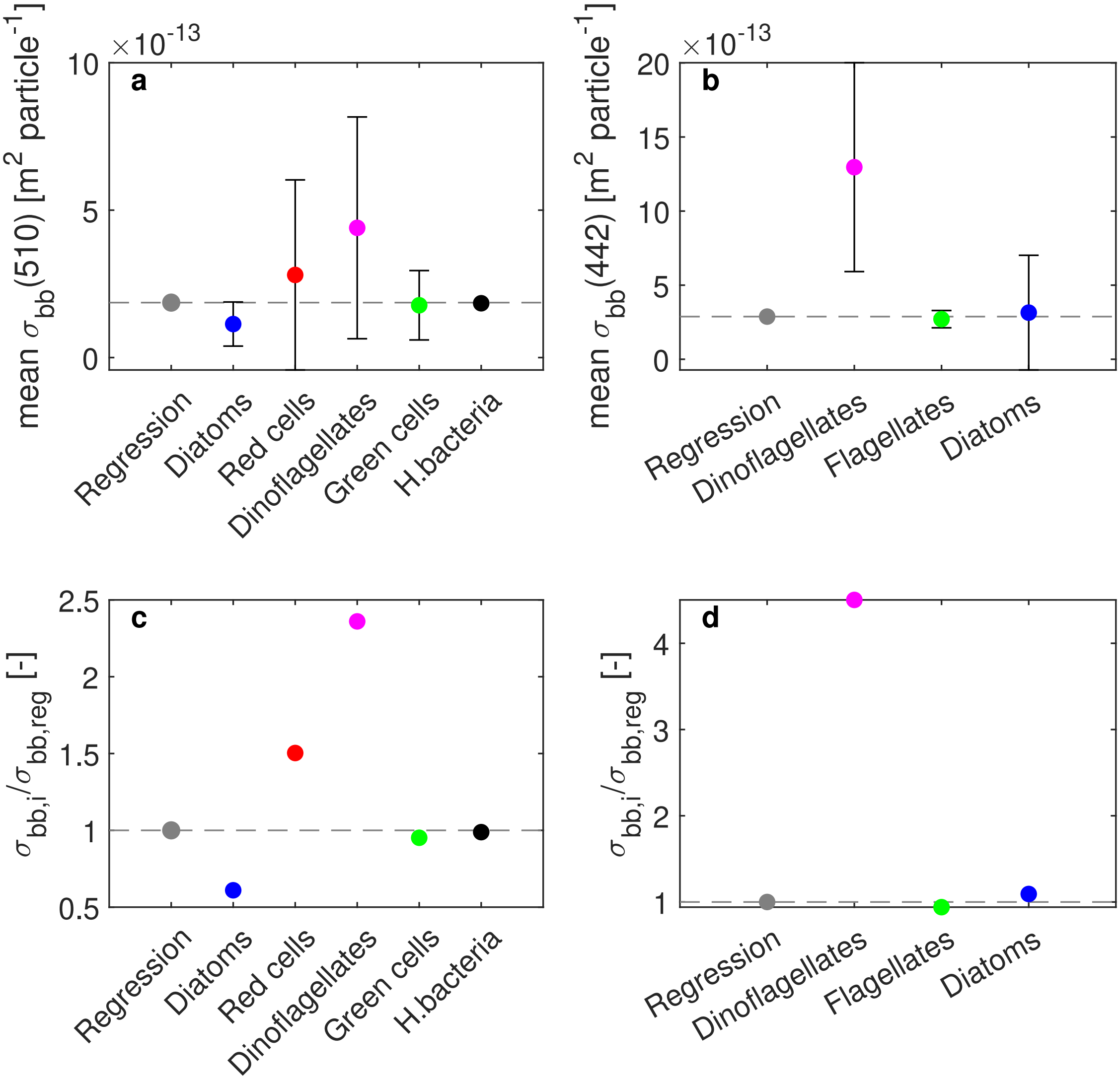


**Figure S5.** (a,b) Mean of $\sigma_{bb}$ for each group after size-corrections from the data sets in Vaillancourt et al. (2004) (left panels) and Whitmire et al. (2010) (right panels) as shown in figure S4 (error-bars show the standard deviation). (c,d) Mean of each group normalized to the value from the regression equation. Before averaging, all data have been corrected for an equivalent diameter of 5$\mu$m using the regression equations $\sigma_{bb,vail}(510) = 4 \times 10^{-15} D^{2.39}$ and $\sigma_{bb,whit}(442) = 10^{-14.48} D^{2.78}$.

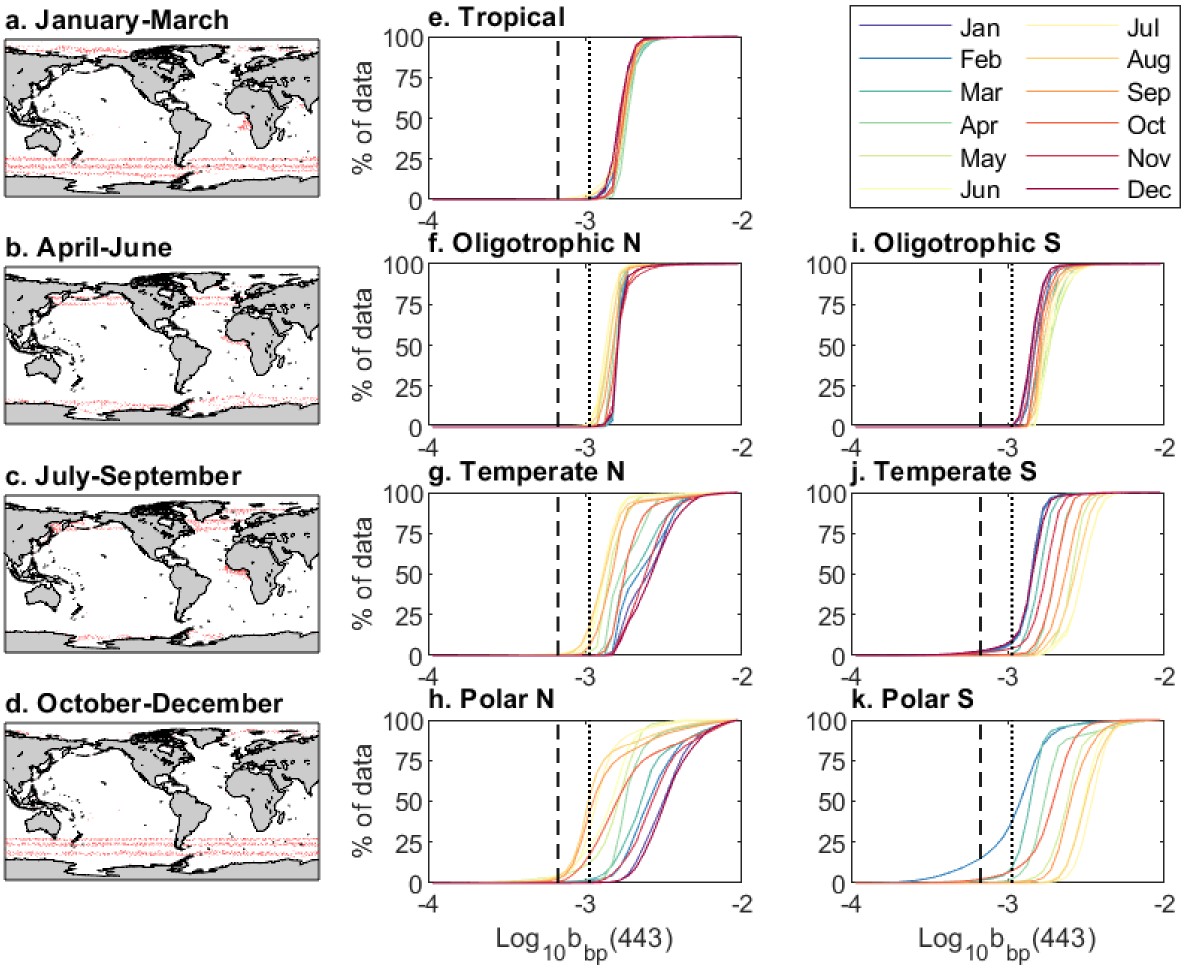


**Figure S6.** (a-d) Data points from satellite remote-sensing (MODIS-GIOP) that are below the threshold $b_{b,crit}$. (e-k) cumulative distributions of all satellite $b_{bp}$ by biomes in the Northern hemisphere (f-h) and southern hemisphere (i-k). Dotted line is $b_{b,crit}$ (where algorithms differ by more than a factor of 3), dashed line is $b_{b,crit2}$ (where algorithms differ by more than an order of magnitude).

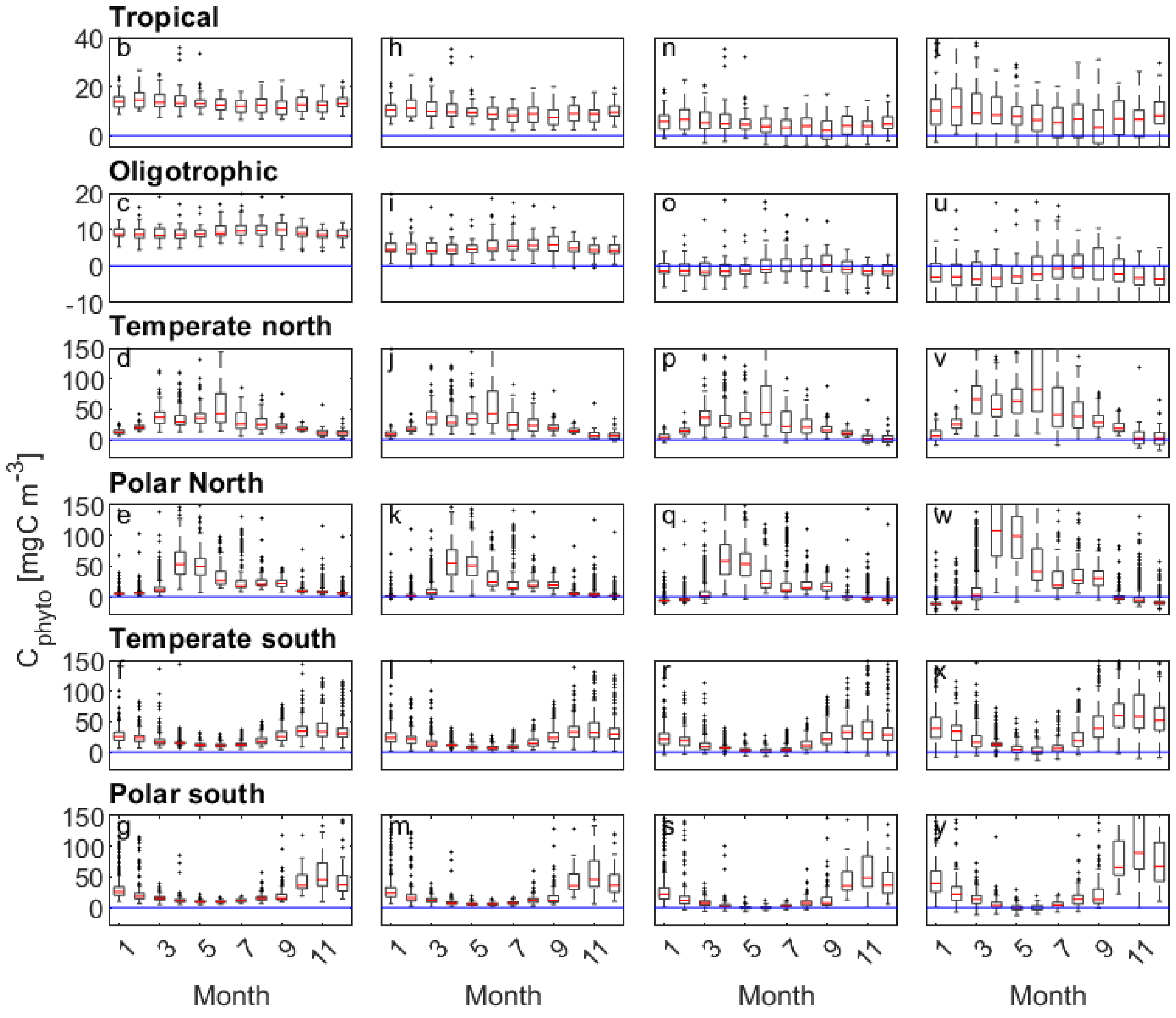


**Figure S7.** $C_{phyto}$ using four different algorithms (columns) using the Bgc-Argo $b_{bp}$ data for different regions. Algorithms are: Graff et al. (2015) (1st column), Behrenfeld et al. (2005) (2nd column), Qiu et al. (2021) (3rd column), and Martinez-Vicente et al. (2013) (4th column).

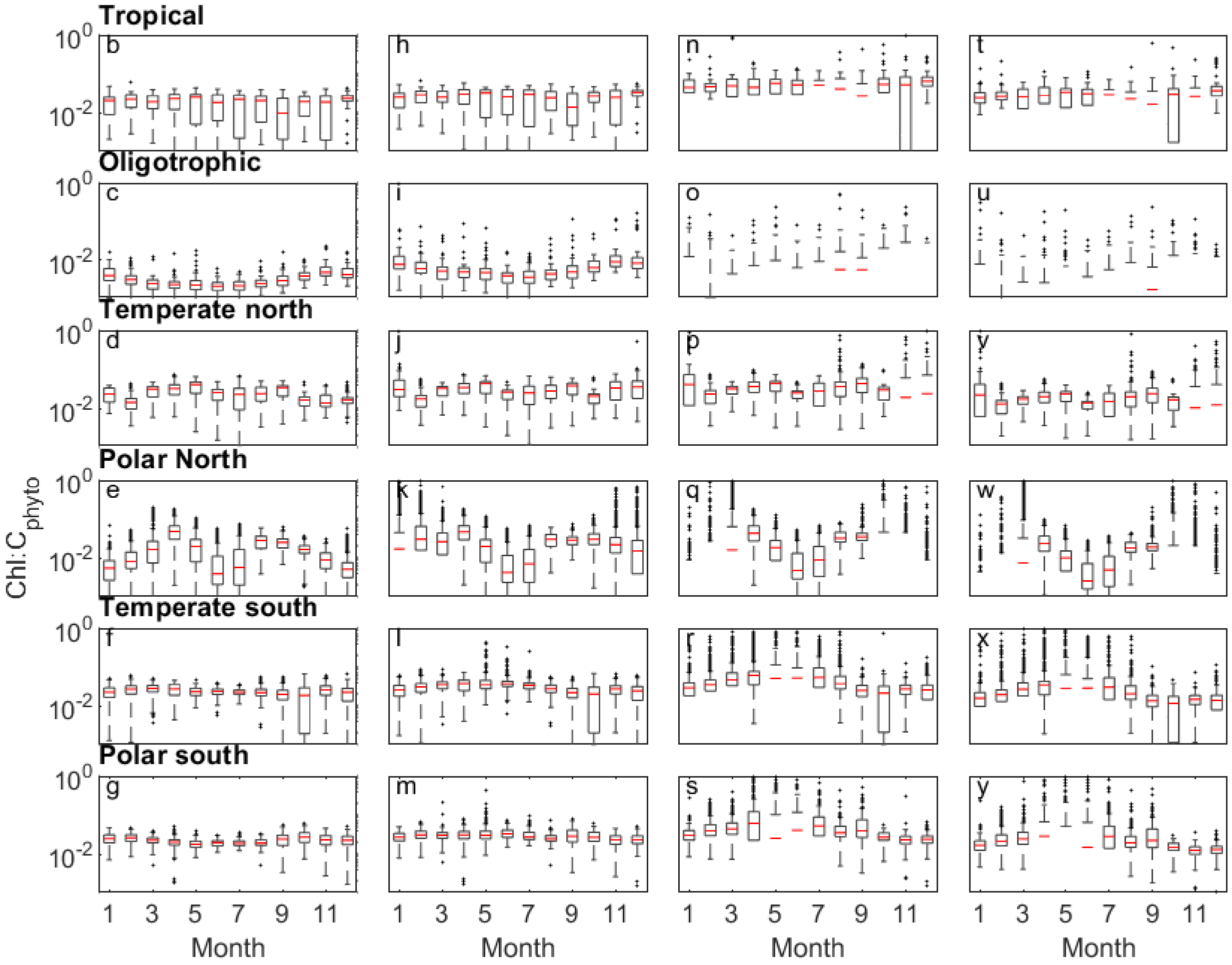


**Figure S8.** $Chl : C_{phyto}$ ratios using four different algorithms (columns) using the Bgc-Argo $b_{bp}$ data for different regions. Algorithms are: Graff et al. (2015) (1st column), Behrenfeld et al. (2005) (2nd column), Qiu et al. (2021) (3rd column), and Martinez-Vicente et al. (2013) (4th column).

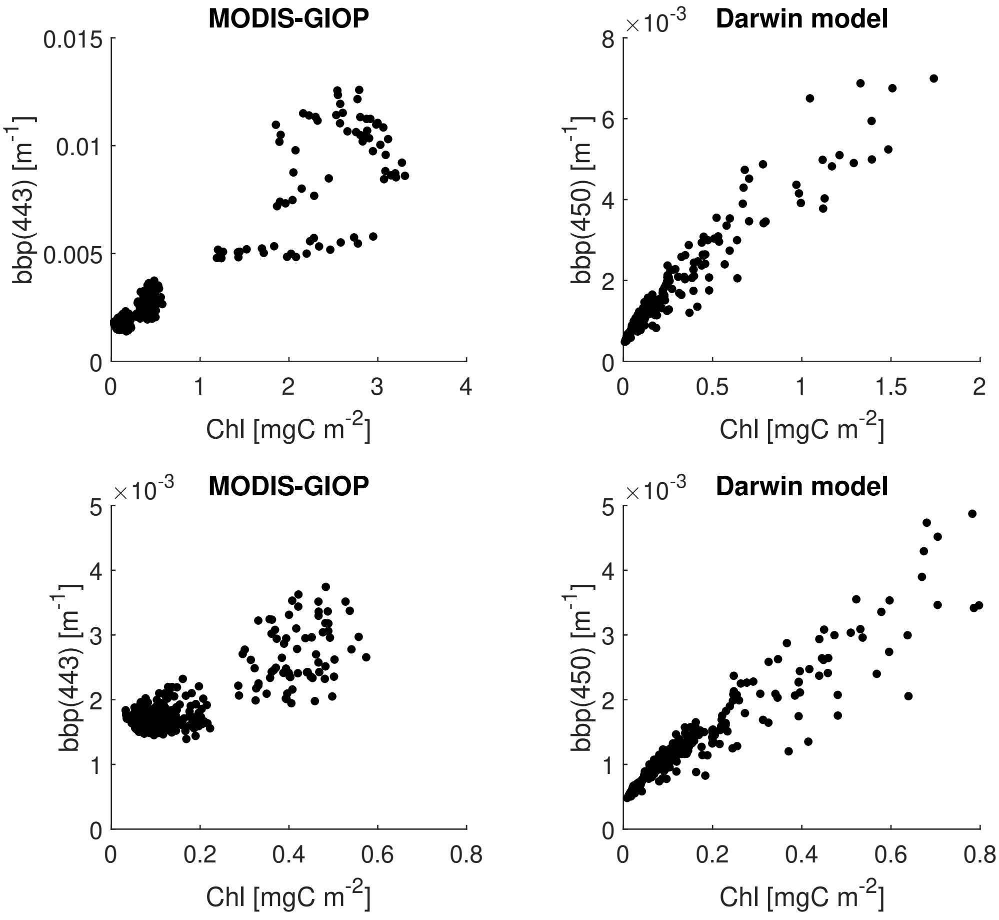


**Figure S9.** (a) Remote sensing (MODIS-GIOP) averaged following the same grouping procedure as in Behrenfeld et al. (2005) and (b) MITgcmBgc model following the same procedure as well. The procedure divides regions according to ranges of the standard deviation of Chl in each bin. Lower panels show the same plot but with the axis constrained to the same ones as in Behrenfeld et al. (2005). Note that the averaging is not exactly the same as in Behrenfeld et al. (2005) because we have used climatological monthly averages (between 2003 and 2021), and not a range of years as in Behrenfeld et al. (2005).

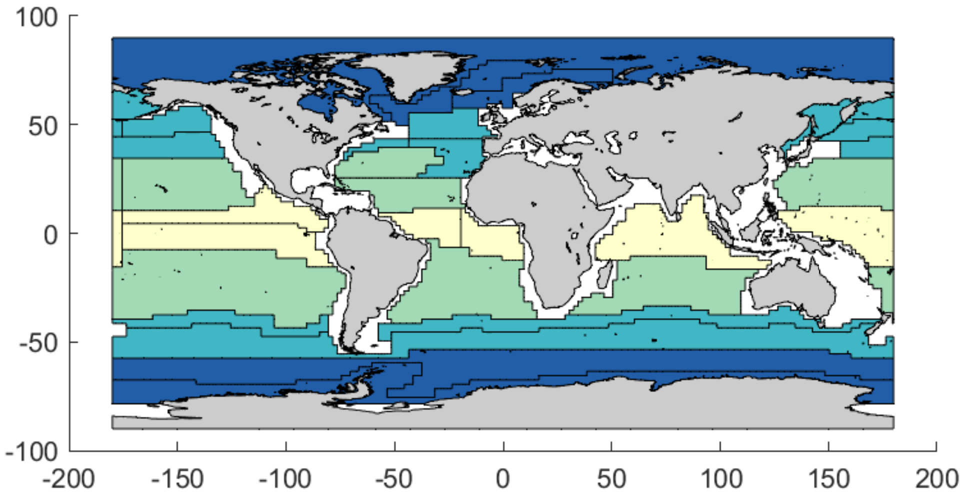


**Figure S10.** Black lines show Longhurst regions. Shades of blue an yellow shoe the regions that we have used in this study. These Longhurst regions have been grouped into four biomes: tropical (yellow), oligotrophic (green), temperate (light blue), sub-polar and polar (dark blue).

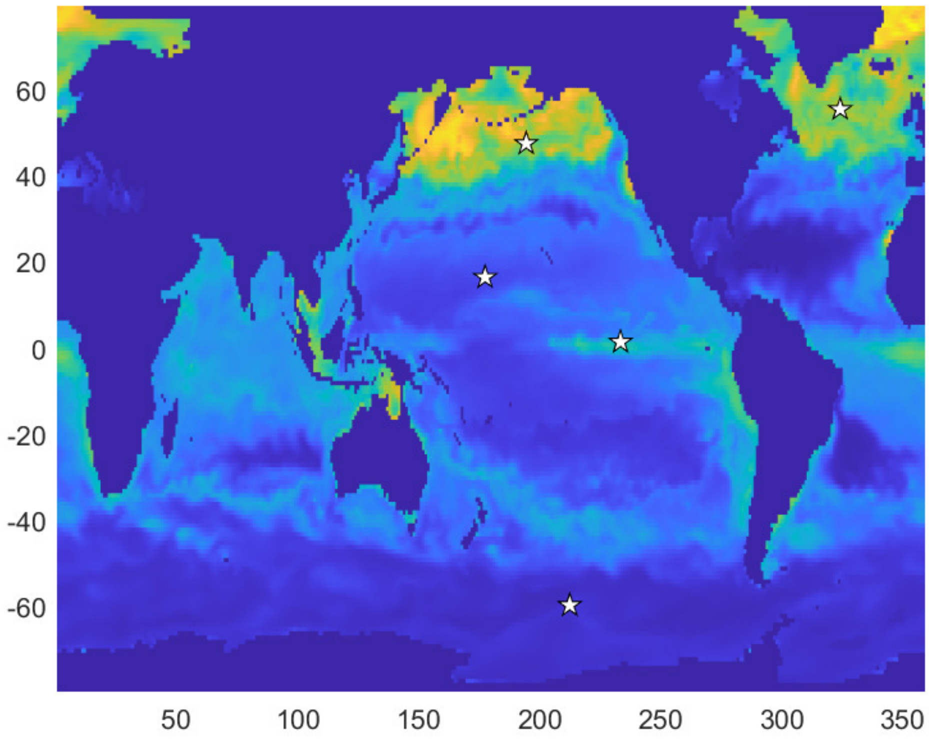


**Figure S11.** Stars show the locations used in figure 7 of the main text.

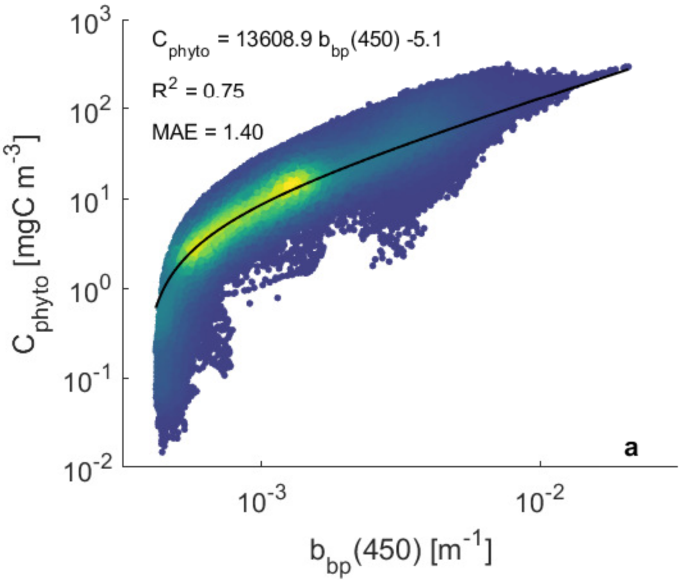


**Figure S12.** Regression assuming that $C_{phyto}$ are only pure autotrophs (without mixotroph biomass included).

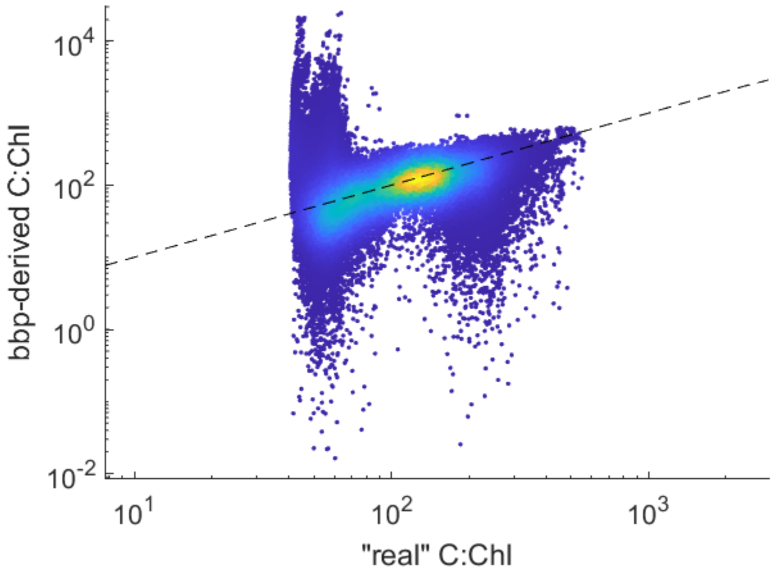


**Figure S13.** $C_{phyto}$:Chl ratio obtained using a bbp-derived $C_{phyto}$ algorithm vs. the "real" $C_{phyto}$:Chl in the model. Dashed line is the 1:1 line.
